# Multiplexed DNA-PAINT Imaging of the Heterogeneity of Late Endosome/Lysosome Protein Composition

**DOI:** 10.1101/2024.03.18.585634

**Authors:** Charles Bond, Siewert Hugelier, Jiazheng Xing, Elena M. Sorokina, Melike Lakadamyali

## Abstract

Late endosomes/lysosomes (LELs) are crucial for numerous physiological processes and their dysfunction is linked to many diseases. Proteomic analyses have identified hundreds of LEL proteins, however, whether these proteins are uniformly present on each LEL, or if there are cell-type dependent LEL sub-populations with unique protein compositions is unclear. We employed a quantitative, multiplexed DNA-PAINT super-resolution approach to examine the distribution of six key LEL proteins (LAMP1, LAMP2, CD63, TMEM192, NPC1 and LAMTOR4) on individual LELs. While LAMP1 and LAMP2 were abundant across LELs, marking a common population, most analyzed proteins were associated with specific LEL subpopulations. Our multiplexed imaging approach identified up to eight different LEL subpopulations based on their unique membrane protein composition. Additionally, our analysis of the spatial relationships between these subpopulations and mitochondria revealed a cell-type specific tendency for NPC1-positive LELs to be closely positioned to mitochondria. Our approach will be broadly applicable to determining organelle heterogeneity with single organelle resolution in many biological contexts.

**Summary:** This study develops a multiplexed and quantitative DNA-PAINT super-resolution imaging pipeline to investigate the distribution of late endosomal/lysosomal (LEL) proteins across individual LELs, revealing cell-type specific LEL sub-populations with unique protein compositions, offering insights into organelle heterogeneity at single-organelle resolution.

## Introduction

The endosomal-lysosomal system is a complex and dynamic network of membrane-bound compartments that plays a critical role in maintaining cellular homeostasis (Klumperman and Raposo, 2014; van Meel and Klumperman, 2008). These compartments include early, recycling and late endosomes, lysosomes and autophagosomes, which are distinguished by their unique functions, sub-cellular locations, sizes, morphologies, and specific luminal and membrane protein compositions (Klumperman and Raposo, 2014; van Meel and Klumperman, 2008). Lysosomes, notable for their acidic lumen containing an array of degradative enzymes (De Duve et al., 1955), mature from late endosomes and represent the final stage in the endosomal-lysosomal pathway (Ballabio and Bonifacino, 2020; Bonifacino and Traub, 2003; Yang and Wang, 2021). Their acidic environment, maintained by the v-ATPase proton pump, is ideal for the activity of lysosomal hydrolases (Mindell, 2012). Historically viewed as the cell’s waste disposal system, lysosomes break down various macromolecules and organelles. However, our current understanding has significantly advanced beyond this historical view, revealing lysosomes as much more dynamic organelles that regulate nutrient sensing, metabolic signaling, membrane repair and several other cellular processes (Lawrence and Zoncu, 2019; Reddy et al., 2001; Settembre et al., 2013; Trivedi et al., 2020) . Additionally, lysosomes are increasingly implicated in numerous neurodegenerative diseases and the aging process (Chen et al., 2019; Malik et al., 2019; Settembre et al., 2013; Tan and Finkel, 2023; Udayar et al., 2022). Given their emerging importance beyond the canonical degradative function, a more in-depth knowledge of lysosomal homeostasis will enable a better understanding of their vital roles in both health and disease.

In electron microscopy studies lysosomes appear as electron dense, spherical organelles with diameters ranging between 200 nm to 600 nm (Barral et al., 2022). While electron microscopy is effective in characterizing the ultrastructure of endosomal/lysosomal compartments, it cannot reveal their protein composition and dynamic behavior. Consequently, light microscopy is indispensable for studying the dynamic interconversion of endosomal/lysosomal compartments, their motility, sub-cellular positioning, and function. These studies typically use fluorescent markers to track the various stages of endosomal/lysosomal compartments. Specifically for lysosomes, probes like lysotracker are employed, which accumulate in organelles with low pH levels (Barral et al., 2022). However, these pH dependent fluorescent dyes often fail to discriminate between late endosomes and lysosomes, as both have acidic luminal pH (Barral et al., 2022). To overcome this limitation, surface markers, which highlight the stage-specific molecular machineries present on the membranes of these compartments, offer a more direct method of discrimination (Klumperman and Raposo, 2014; van Meel and Klumperman, 2008). For instance, membrane proteins such as EEA1 and Rab5 are indicative of early endosomes, while Rab7 is a marker for late endosomes (Lakadamyali et al., 2006; Nielsen et al., 1999; Rink et al., 2005; Vanlandingham and Ceresa, 2009; Wilson et al., 2000).

The most abundant lysosomal membrane proteins are the lysosome associated membrane proteins 1 (Lippincott-Schwartz and Fambrough, 1986) and 2 (LAMP1 and LAMP2), lysosomal integral membrane protein 2 (LIMP2) and CD63 (LIMP1/LAMP3) (Lubke et al., 2009; Schroder et al., 2010; Schwake et al., 2013; Winchester, 2001). LAMP1/2 have been suggested to play roles in lysosome biogenesis (Schwake et al., 2013) and in regulating lysosomal pH through direct interactions with the channel TMEM175 (Zhang et al., 2023). CD63, a member of the tetraspanin superfamily is upregulated in many cancers (Pols and Klumperman, 2009) and may play roles in extracellular vesicle production and endosomal cargo sorting (Hurwitz et al., 2018; van Niel et al., 2011). Due to their high abundance, LAMP1 and LAMP2 are commonly used as surface markers in light microscopy studies of lysosomes. These proteins are often tagged with a fluorescent protein and over-expressed for visualization during live or fixed cell imaging to study lysosomal dynamics and functions. However, the over-expression of these markers might alter the dynamics, distribution, pH and functionality of the endosomal-lysosomal compartments.

Beyond LAMP and LIMP proteins, proteomic studies have identified over 100 different lysosomal membrane proteins, including ion channels, transporters, and exchangers (Akter et al., 2023; Bagshaw et al., 2005; Lubke et al., 2009; Muthukottiappan and Winter, 2021; Schroder et al., 2010; Yu et al., 2024). The lysosomal membrane also serves as a hub for various proteins that dynamically and transiently assemble on it including components of the nutrient sensing mTORC pathway such as mTORC1, Ragulator, and Raptor (Perera and Zoncu, 2016; Rogala et al., 2019; Sancak et al., 2010; Sancak et al., 2008; Settembre et al., 2013; Zoncu et al., 2011).

A key question remains: are all these proteins identified in proteomic studies equally abundant on every lysosome? Considering the heterogeneity and continuous exchange between endosomes and lysosomes, there could be a continuous gradient of organelles that all have the same protein complement but in varying levels of abundance. It is also plausible that there are molecularly distinct organelles, each containing a unique complement of lysosomal proteins. Addressing this key question requires a method that can visualize and quantify a large number of lysosome-associated proteins in a multiplexed fashion with high molecular specificity, sensitivity and spatial resolution. High spatial resolution is particularly important in the context of resolving the small, densely packed lysosomes within cells. Previous work used correlative light and electron microscopy to reveal differences in the molecular composition of early and late endosomes (van der Beek et al., 2022). However, this approach is limited in its scope as it is low-throughput and technically challenging. In addition, the reliance on low resolution conventional microscopy limits it to evaluating sub-cellular compartments that are spatially well-separated within the cell. Finally, conventional immunofluorescence microscopy is not sensitive towards detecting low abundance proteins and is not highly quantitative.

Super-resolution light microscopy has emerged as a powerful tool for visualizing the inner architecture of cells with nanoscale spatial resolution (Bond et al., 2022). Among various super-resolution methods, DNA Point Accumulation in Nanoscale Topography (DNA-PAINT) stands out for its ability to multiplex (Jungmann et al., 2014). Multiplexed DNA-PAINT or Exchange-PAINT employs DNA-barcoded antibodies to detect and sequentially image multiple proteins (Jungmann et al., 2014). DNA-PAINT’s single molecule detection efficiency also makes it highly sensitive, allowing for the detection of even low-abundance proteins. Importantly, DNA-PAINT is highly quantitative. The well-defined binding and unbinding kinetics of the imager oligonucleotides ensure that the number of detected localizations is directly and linearly proportional to the abundance of the targeted protein (Jungmann et al., 2016). This quantitative aspect of DNA-PAINT makes it ideally suited for accurate analysis of protein levels across different endosomal and lysosomal compartments.

Here, we developed a quantitative pipeline using multiplexed DNA-PAINT imaging to analyze the protein abundance and heterogeneity on late endosomal and lysosomal membranes at the endogenous level. This was achieved using thoroughly validated antibodies in two different cell types, HeLa and ARPE-19 cells. Our findings reveal that the canonical and abundant lysosomal proteins, LAMP1 and LAMP2, mark the same population of organelles. Therefore, we used LAMP1 as a reference to determine the abundance of other lysosomal proteins in these LAMP1-positive compartments, which we refer to as late endosomes/lysosomes or LELs for short. Interestingly, CD63, another canonical and abundant lysosomal protein, exhibited more variation, especially in ARPE-19 cells, where it appeared only on a subset of LELs. Niemann Pick Disease Type C1 protein (NPC1), which plays a role in cholesterol trafficking on the lysosomal membrane (Infante et al., 2008; Pfeffer, 2019), formed small nanodomains that were present only on distinct LEL subsets in both cell types suggesting that only a specific subset of LELs is engaged in NPC1-mediated cholesterol homeostasis. LAMTOR4, a subunit of the Ragulator complex involved in mTOR activation (Sancak et al., 2010; Sancak et al., 2008; Zoncu et al., 2011), formed larger nanodomains compared to NPC1 outside the membrane of a subset of LELs. 4-color multiplexed imaging revealed several LEL subpopulations, each characterized by their unique combination of membrane proteins, showing diverse, molecularly distinct LEL subsets. Spatial analysis also provided insights into the subcellular localization of these distinct subsets, particularly in relation to the nucleus and other organelles, such as mitochondria.

Overall, our study offers quantitative tools and a novel framework for characterizing the protein composition of individual organelles within cells with high sensitivity and spatial resolution, revealing the heterogeneity of LELs characterized by both unique combination of resident proteins and variability in their abundance. This method can be widely applied to investigate organelle heterogeneity in various cellular contexts.

## Results

### Quantitative DNA-PAINT pipeline for characterizing LEL membrane proteins

We first developed a comprehensive quantitative pipeline designed for broad application in visualizing and quantifying the abundance of various proteins on organelle membranes and applied it to characterize the abundance of six LEL proteins: LAMP1, LAMP2, CD63, TMEM192, NPC1 and LAMTOR4 (**Figure 1**). Since we aimed to characterize LEL proteins at their endogenous levels, we used immunofluorescence (IF) labeling with commercially available antibodies that have previously been validated in IF studies (Cason et al., 2022; Eapen et al., 2021; Gallagher and Holzbaur, 2023; Hiragi et al., 2022; Keren-Kaplan et al., 2022; Rebsamen et al., 2015; Wang et al., 2020b; Weng et al., 2022). To further validate antibody specificity, we over-expressed target proteins fused to a tag (when available) and compared the antibody staining to that of the tag (**Supplementary Figure 1 A-E**). In all cases, we observed a high degree of co-localization between the tagged protein and the antibody stain on vesicular compartments (**Supplementary Figure 1 A-E**), validating the specificity of the used antibodies. We additionally validated antibodies against several low abundance proteins with knockdown and knockout cell lines (see below for more details).

**Figure 1:**
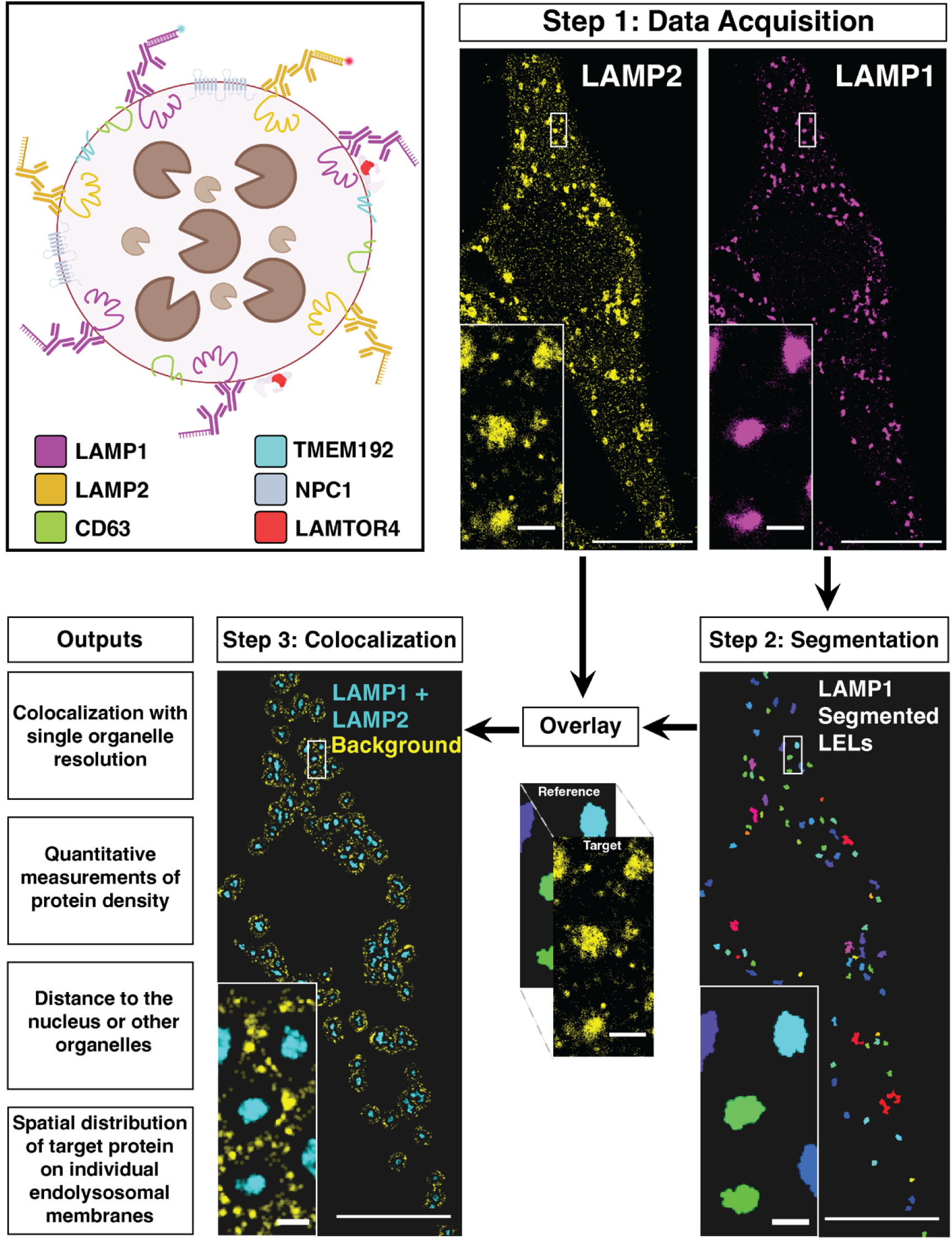
Quantitative DNA-PAINT pipeline for characterizing LEL membrane proteins. Schematic shows LELs with target proteins imaged in this study. Target proteins (e.g. LAMP1 and LAMP2) are labeled with primary and secondary antibodies for DNA-PAINT imaging. Step 1 shows the data acquisition step in which a dual-color DNA-PAINT image is acquired (in this case LAMP1 and LAMP2). Subsequently, in Step 2, LAMP1 is segmented to be used as the reference channel, and the LAMP2 raw localizations (target channel) are overlaid onto the segmented LELs. During co-localization (Step 3), the target localization density is measured in the region of the segmented LELs (co-localization region), as well as a local background region surrounding the LEL. If the density within the co-localization region is significantly (more than 3 standard deviations) enriched over the background density, that LEL is determined to have both target and reference proteins. Several quantitative outputs are obtained as a result of this pipeline. Cell scale bars = 10 µm, inset scale bars = 1 µm.

Given the high abundance of LAMP1 and LAMP2 on LEL membranes, we validated our quantitative pipeline using DNA-PAINT imaging of these two proteins in two distinct cell types: HeLa (**Figure 1 and 2A**) and ARPE-19 (**Supplementary Figure 2A**). As expected, both proteins predominantly localized to vesicular compartments, resembling LEL structures. We first optimized fixation and permeabilization by comparing two different methods aiming to minimize any potential disruption to the LEL localization of LAMP proteins. Previous studies showed that fixation with -20°C methanol disrupts cell and organelle membranes and is not suitable for visualizing membrane bound organelles (Whelan and Bell, 2015). We therefore refrained from using this specific fixation and instead evaluated the conventional aldehyde-based fixation using 4% Paraformaldehyde (PFA) against Glyoxal fixation (**Supplementary Figure 2F**), the latter suggested to be a quicker fixative (Richter et al., 2018) (see Methods). In addition, we compared the standard Triton-X100 permeabilization with the gentler Saponin (**Supplementary Figure 2G-I**) (see Methods). Our results indicated that a combination of 4% PFA with Saponin most effectively maintained the vesicular enrichment of LAMP1 and LAMP2 (**Supplementary Figure 2G-I**). Consequently, we adopted this combination for all subsequent experiments and analyses.

**Figure 2:**
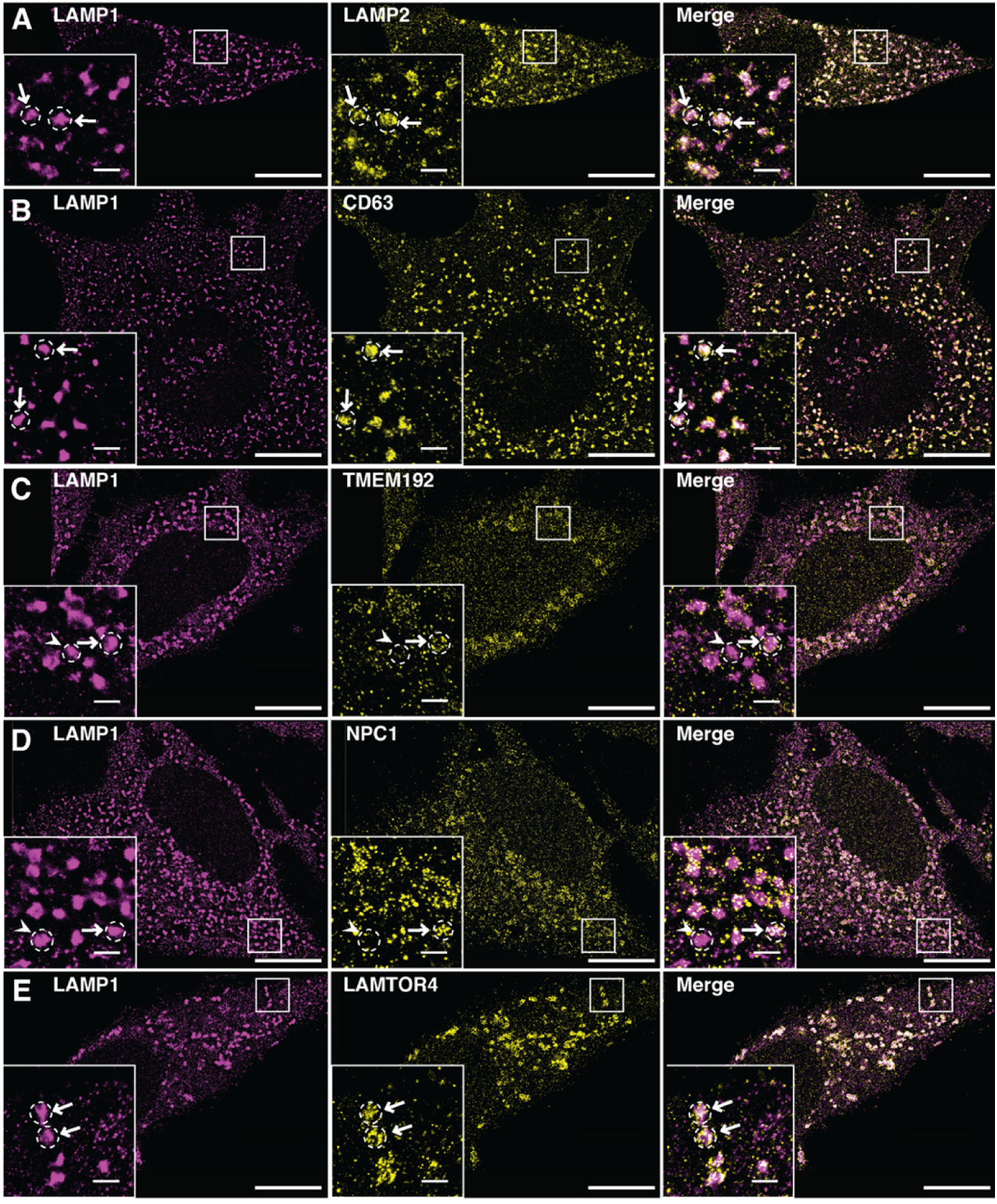
Dual-Color DNA-PAINT in HeLa cells identifies heterogeneity in the enrichment of canonical LEL proteins. Representative DNA-PAINT images of LAMP1 reference channel (magenta) and target protein channel (yellow) in HeLa cells for **A.** LAMP2 **B.** CD63 **C.** TMEM192 **D.** NPC1 and **E.** LAMTOR4. Arrows indicate LELs positive for both LAMP1 and target protein, arrowheads indicate LELs positive for LAMP1 and negative for target protein, dotted circle indicates which LEL arrow or arrowhead refers to. Cell scale bars, 10 µm. Inset scale bars, 1 µm.

We next developed a robust co-localization analysis to determine the extent of co-localization between LAMP1 and LAMP2-positive LELs in dual-color DNA-PAINT images. While several co-localization algorithms for super-resolution microscopy exist, they predominantly rely on cross-correlation of point localizations (Hugelier et al., 2023; Malkusch et al., 2012; McCall, 2024; Stone and Veatch, 2015). This approach is not applicable to quantifying object-based co-localization typical of LELs. To address this, we developed a robust, object-based co-localization algorithm (see Methods for details). To this end, we first segmented individual LELs in a reference channel (e.g. LAMP1, Ch_REF_) using Voronoi-based clustering and segmentation (**Figure 1**). Next, we applied these segmented compartments as a mask to compute the density of localizations from the second, target channel (Ch_TARGET_) within the masked area (**Figure 1**). Furthermore, we calculated a background density, representative of the local background density proximal to the mask (see Methods for details) (**Figure 1**). When the localization density inside the reference mask exceeded three standard deviations above the background density, we interpreted this as a significant signal surpassing background levels, indicative of positive co-localization.

This analysis enabled us to determine both the percentage of LAMP1-positive LELs that exhibited co-localization with LAMP2-positive LELs, as well as the density of LAMP2 protein within these LAMP1-positive LELs (**Figures 3A-C**). Recognizing that in DNA-PAINT, localization density correlates with imaging duration, we standardized this duration across all proteins imaged to guarantee both reproducibility and consistent comparison. We also verified that the imaging time was adequately long to capture majority of the localizations associated with the protein being imaged. To this end, we computed the coverage area of target protein localizations per LEL within the reference mask. While the coverage percentage varied between individual LELs depending on the abundance of the target protein, we observed levels approaching and reaching saturation for both LAMP1 and LAMP2 for each LEL (**Supplementary Figure 3A, B**), confirming that the imaging duration was sufficient for capturing majority of relevant protein localizations within the mask. Overall, these results support that our quantitative imaging and analysis pipeline is capable of reliably determining protein density within LEL compartments.

**Figure 3:**
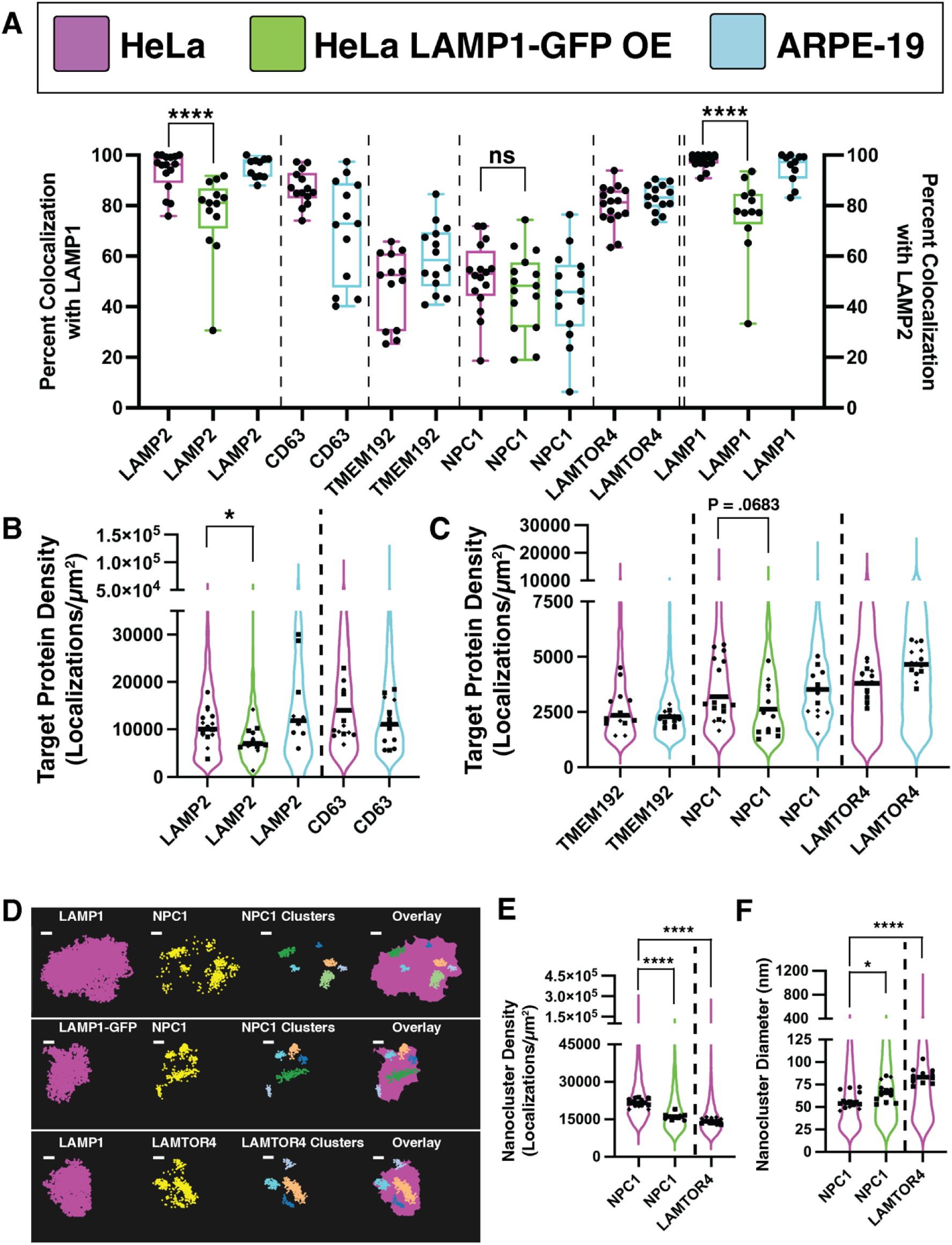
Dual-color DNA-PAINT identifies LEL subpopulations with variable protein makeup that persist across epithelial cell types. **A.** Box and whisker plots showing the percent co-localization of target proteins with reference LELs in HeLa, HeLa LAMP1-GFP overexpressing, or ARPE-19 cells. All targets were imaged in three independent biological replicates. LAMP1 was used as the reference for LAMP2 (n = 16 HeLa cells, 13 LAMP1-GFP overexpressing cells, 11 ARPE-19 cells), CD63 (14 HeLa cells, 13 ARPE-19 cells), TMEM192 (13 HeLa cells, 14 ARPE-19 cells), NPC1 (n = 16 HeLa cells, 15 LAMP1-GFP overexpressing cells, 14 ARPE-19 cells), and LAMTOR4 (16 HeLa cells, 14 ARPE-19 cells) (left Y-axis), while LAMP2 was used as the reference for LAMP1 (n = 16 HeLa cells, 13 LAMP1-GFP overexpressing cells, 11 ARPE-19 cells) (right Y-axis). Plot line color indicates cell type, black circles indicate individual cells. Mann-Whitney U test was performed to compare percent co-localization of LAMP2 with LAMP1 (n = 16 HeLa cells, n = 13 HeLa LAMP1-GFP OE cells) (P < 0.0001), NPC1 with LAMP1 (n = 16 HeLa cells, n = 13 HeLa LAMP1-GFP OE cells) (P = 0.3527, no significance), and LAMP1 with LAMP2 (n = 16 HeLa cells, n = 12 HeLa LAMP1-GFP OE cells) (P < 0.0001) in HeLa vs. HeLa LAMP1-GFP overexpressing (OE) cells. **B-C**: Violin plots of target protein density for high density **(B)** and low density **(C)** targets. Plot line color indicates cell type, black line indicates median target density on LELs positive for a given target. Black circles, squares, and diamonds represent individual cells from 3 different biological replicates. Mann-Whitney U test was performed to compare median target densities in HeLa vs HeLa LAMP1-GFP overexpressing cells for **B.** LAMP2 (n = 16 HeLa cells, n = 13 HeLa LAMP1-GFP OE cells) (P = 0.0172) and **C.** NPC1 (n = 16 HeLa cells, n = 13 HeLa LAMP1-GFP OE cells) (P = 0.0683). **D-F:** Pattern analysis of target proteins on individual LEL membranes. **(D)** Representative zoomed-in DNA-PAINT image of an individual LEL as defined by LAMP1 or LAMP1-GFP localizations (magenta, leftmost panel), raw localizations of target protein on the LEL (yellow, second panel), nanoclusters of the target channel segmented using DBScan and pseudocolored to indicate distinct nanoclusters (third panel) and these nanoclusters overlaid on LEL reference channel alphashape bounding area calculated from the LAMP1 or LAMP1-GFP localizations. Violin plots **(E)** show nanocluster density (number of localizations of the target protein per unit area of the nanocluster). Plot line color indicates cell type, black line indicates median nanocluster density on LELs positive for a given target. Black circles, squares, and diamonds represent individual cells from 3 different biological replicates. Mann-Whitney U test was performed to compare median nanocluster densities in HeLa cells between NPC1 (n = 16 cells) and LAMTOR4 (n = 16 cells) (P <0.0001) and for NPC1 in HeLa (n =16 cells) vs HeLa LAMP1-GFP OE cells (n = 13 cells) (P <0.0001). Violin plots **(F)** show nanocluster diameter. Mann-Whitney U test was performed to compare median nanocluster diameters in HeLa cells between NPC1 (n = 16 cells) and LAMTOR4 (n = 16 cells) (P <0.0001) and for NPC1 in HeLa (n =16 cells) vs HeLa LAMP1-GFP OE cells (n = 13 cells) (P <0.0251).

The co-localization analysis revealed 93.5±7.8% and 95.0±4.0% overlap between LAMP1-positive LELs and LAMP2-positive ones both in HeLa and ARPE-19 cells (**Figure 3A**), respectively, as expected. This result held true when LAMP2 was used as a reference mask instead of LAMP1 (97.7±2.7% overlap for HeLa, 94.9±6.0% for ARPE-19 cells) (**Figure 3A**), which underscores the robustness of our analysis pipeline. Additionally, as a negative control, the early endosomal marker EEA1 displayed minimal co-localization with LAMP1 (14.1±6.1% overlap for HeLa, 10.7±5.1% for ARPE-19 cells) (**Supplementary Figure 3C, D**), further validating the co-localization analysis pipeline.

The immunofluorescence labeling of two proteins positioned in close proximity on the LEL membrane might be impeded by steric effects. To test this, we determined the localization density of LAMP1 and LAMP2 on each LEL using our pipeline. If steric effects were present, an inverse correlation would be expected, where high density of one protein would coincide with low density of the other. However, we observed low correlation between the densities of LAMP1 and LAMP2 proteins on LELs (**Supplementary Figure 3E, F**), ruling out concerns of labeling-related steric interference.

Overall, we have successfully developed a robust and quantitative DNA-PAINT imaging pipeline, enabling precise determination of protein levels and their co-localization within LELs.

### Dual-color DNA-PAINT identifies LEL subpopulations with variable protein makeup that persist across epithelial cell types

We applied our quantitative pipeline to determine whether four additional LEL associated proteins, crucial for different aspects of LEL biology, are ubiquitously present on all LAMP1-positive compartments or if there exist subpopulations of LELs with diverse membrane protein compositions (**Figures 2 B-E, Figure 3 A-C, Supplementary Figure 2B-E**). For each protein, we first confirmed that the imaging duration was sufficiently long to capture the majority of the protein localizations within the reference LAMP1 mask as previously described (**Supplementary Figure 3A, B**).

We found that CD63 (also known as LIMP1 or LAMP3), another prevalent LEL protein (Schwake et al., 2013), was present on 87.0±6.8% of LAMP1-positive LELs in HeLa cells (**Figures 2B and 3A**). Interestingly, in ARPE-19 cells, CD63 displayed more variation in its co-localization with LAMP1, ranging from as low as 40% in some cells to almost complete co-localization in others (**Supplementary Figure 2B and Figure 3A**). This result may reflect differences in the maturity or function of LELs in the two different cell types.

The transmembrane protein 192 (TMEM192) is a lesser-known LEL protein, initially identified through organellar proteomics (Chapel et al., 2013; Nguyen et al., 2017; Schroder et al., 2007; Schroder et al., 2010). Although the function of TMEM192 is unclear (Nguyen et al., 2017), it is widely utilized in Lyso-IP proteomics and metabolomics studies to immunoprecipitate lysosomes, as it preserves lysosomal association when over-expressed (Abu-Remaileh et al., 2017). We aimed to determine the LEL localization of this protein under native conditions. We found that TMEM192, while not highly abundant (**Figure 3C**), was consistently present above background levels on 47.9±14.6% of LAMP1-positive LELs in HeLa cells and 59.4±13.1% in ARPE-19 cells (**Figures 2C and 3A, Supplementary Figure 2C**). This result was also corroborated using a second, alternative TMEM192 antibody (**Supplementary Figure 4A, F**). When TMEM192 was knocked down (**Supplementary Figure 4B, E**), there was a marked reduction in both the co-localization percentage and protein density on LELs (**Supplementary Figure 4B, F, G**). Conversely, over-expression of TMEM192 led to a significant increase in protein density and co-localization (81.1±26.0%), consistent with expectations for LEL localization upon over-expression (**Supplementary Figure 4C, F, G**). These results validate that the TMEM192 antibody is specific and sensitive to TMEM192 levels. Additionally, these results demonstrate the capability of DNA-PAINT imaging and our quantitative pipeline to accurately identify and quantify proteins on LEL membranes, even those that are low in abundance. The increase in LAMP1-positive LELs that are also TMEM192-positive when the protein is over-expressed suggests that the LELs analyzed with TMEM192 over-expression are fundamentally different from those identified through TMEM192 endogenous detection.

Niemann-Pick type C1 (NPC1) is an important LEL membrane protein essential for cholesterol export from LELs, with genetic mutations in NPC1 leading to Niemann-Pick type C disease (Infante et al., 2008; Pfeffer, 2019). Analysis of dual-color DNA-PAINT images showed that NPC1 is also lowly abundant on the membrane of 51.6±14.0% of LELs in HeLa cells and 46.5±15.8% in ARPE-19 cells.(**Figures 2D and 3A, C and Supplementary Figure 2D**). DNA-PAINT imaging in NPC1 knockout (KO) HeLa cells (**Supplementary Figure 4D, E**) once again confirmed the specificity of the antibody and the accuracy of the observed signal (**Supplementary Figure 4D-G**). Notably, NPC1 localizations formed densely and tightly packed nanoscale domains on the LEL membrane in DNA-PAINT images (**Figure 3D-F**). We employed DBSCAN clustering to segment and quantitatively analyze these nanoclusters (**Figure 3D**). Our analysis indicated an average of 5 NPC1 nanoclusters per LEL membrane with a median diameter of 55 nm (**Figure 3F**). These results suggest that NPC1 organizes into multiple nanoscale platforms on LEL membranes, potentially facilitating cholesterol export through clustering. These results further underscore the power of DNA-PAINT super-resolution imaging in uncovering the existence of these nanoscale protein assemblies on organelle membranes.

Finally, LAMTOR4, a component of the pentameric Ragulator complex, plays a pivotal role as a scaffold for Rag GTPases crucial in the recruitment and activation of mTORC1 on LEL membranes (Sancak et al., 2010; Sancak et al., 2008; Zoncu et al., 2011). Similar to TMEM192 and NPC1, LAMTOR4 was found in low abundance on LELs (**Figure 2E**, **Figure 3C and Supplementary Figure 2E**). However, in contrast to these proteins, LAMTOR4 was detected on a much higher percentage of LELs (80.0±8.1% in HeLa, 83.0±5.3% in ARPE-19 cells) (**Figure 3A**). DBSCAN clustering analysis revealed distinct characteristics of LAMTOR4 distribution; compared to NPC1, LAMTOR4 formed larger (83 nm), less dense nanoplatforms on the LEL membrane, indicating a different organizational pattern for this protein (**Figure 3D-F**).

After establishing the baseline abundance of LEL proteins and the heterogeneity in LEL subpopulations, we explored how the over-expression of LAMP1, a common practice in cell biology studies focusing on lysosomes, might influence these baseline populations. We over-expressed LAMP1 fused to GFP and used a GFP nanobody to simultaneously visualize LAMP1 and either LAMP2 or NPC1 in dual-color DNA-PAINT images (**Supplementary Figure 4H, I**). We picked LAMP2 and NPC1 for our analysis as they provide examples of both high and low abundance LEL proteins. Quantitative analysis showed that over-expression differentially affected these two proteins. Notably, the levels of both LAMP2 and NPC1 slightly decreased on LEL membranes following LAMP1 over-expression (**Figure 3B, C**). This decrease was a result of an increase in the size of LELs rather than a decrease in the absolute amount of LAMP1 or NPC1 (**Supplementary Figure 4J**). Interestingly, while the overlap percentage of LAMP1-positive LELs with NPC1 remained unchanged (**Figure 3A**), there was a significant reduction in the co-localization of LAMP1-positive LELs with LAMP2 – decreasing from 93.5±7.8% to 76.8±16.2% (**Figure 3A**). Similar trends were observed when LAMP2 was used as the reference, showing a decrease in the overlap with LAMP1 (**Figure 3A**). Additionally, the over-expression of LAMP1 significantly influenced NPC1 nanoclusters, resulting in an increase in their size and reduction in their packing density on the LEL membrane (**Figure 3E, F**). These results highlight that the over-expression of LEL proteins can affect not only the levels of the over-expressed protein itself, but also the size of the organelle, the density, as well as the organization of other membrane proteins. Moreover, over-expression can lead to LEL subpopulations absent under native conditions such as those that are LAMP1-positive but LAMP2-negative, potentially due to abnormal incorporation of LAMP1 to compartments that normally lack it or emergence of novel LEL sub-populations.

Finally, the dual-color imaging gave us the opportunity to test the robustness and the quantitative accuracy of DNA-PAINT imaging as LAMP1 was independently imaged as a reference channel while concurrently evaluating the other five proteins (LAMP2, CD63, TMEM192, NPC1 and LAMTOR4). When we compared the average protein density of LAMP1 across these five distinct biological replicate experiments in HeLa cells, we found no statistically significant differences in its abundance (**Supplementary Figure 3G**). This consistency highlights the high level of quantitative precision and robustness inherent in our imaging and analysis pipeline.

### Spatial analysis reveals the location of LEL subsets with respect to the nucleus and mitochondria

The spatial positioning of lysosomes within a cell is crucial for their function (Pu et al., 2016) and has been shown to be linked to anabolic and catabolic responses and nutrient availability (Jia and Bonifacino, 2019; Korolchuk et al., 2011; Pu et al., 2017; Walton et al., 2018). These links between lysosome positioning and function led us to investigate whether specific LEL sub-populations, characterized by their protein composition, occupy distinct spatial locations relative to the cell nucleus or other cellular organelles. To explore this, we developed a quantitative method to measure the normalized distance of each LEL from the cell nucleus (**Figure 4A-D, Supplementary Figure 5A-D**), identifiable in our super-resolution images as the empty, dark spaces within the cell (Methods). While there were mild trends such as LAMTOR4 or TMEM192-positive LELs being slightly spatially closer to the cell nucleus in ARPE19 cells, these trends were not statistically significant (**Supplementary Figure 5B-D**). These findings suggest that, under homeostatic conditions, different subpopulations of LELs are distributed across a wide range of spatial locations, extending from the perinuclear area to the cell periphery.

**Figure 4:**
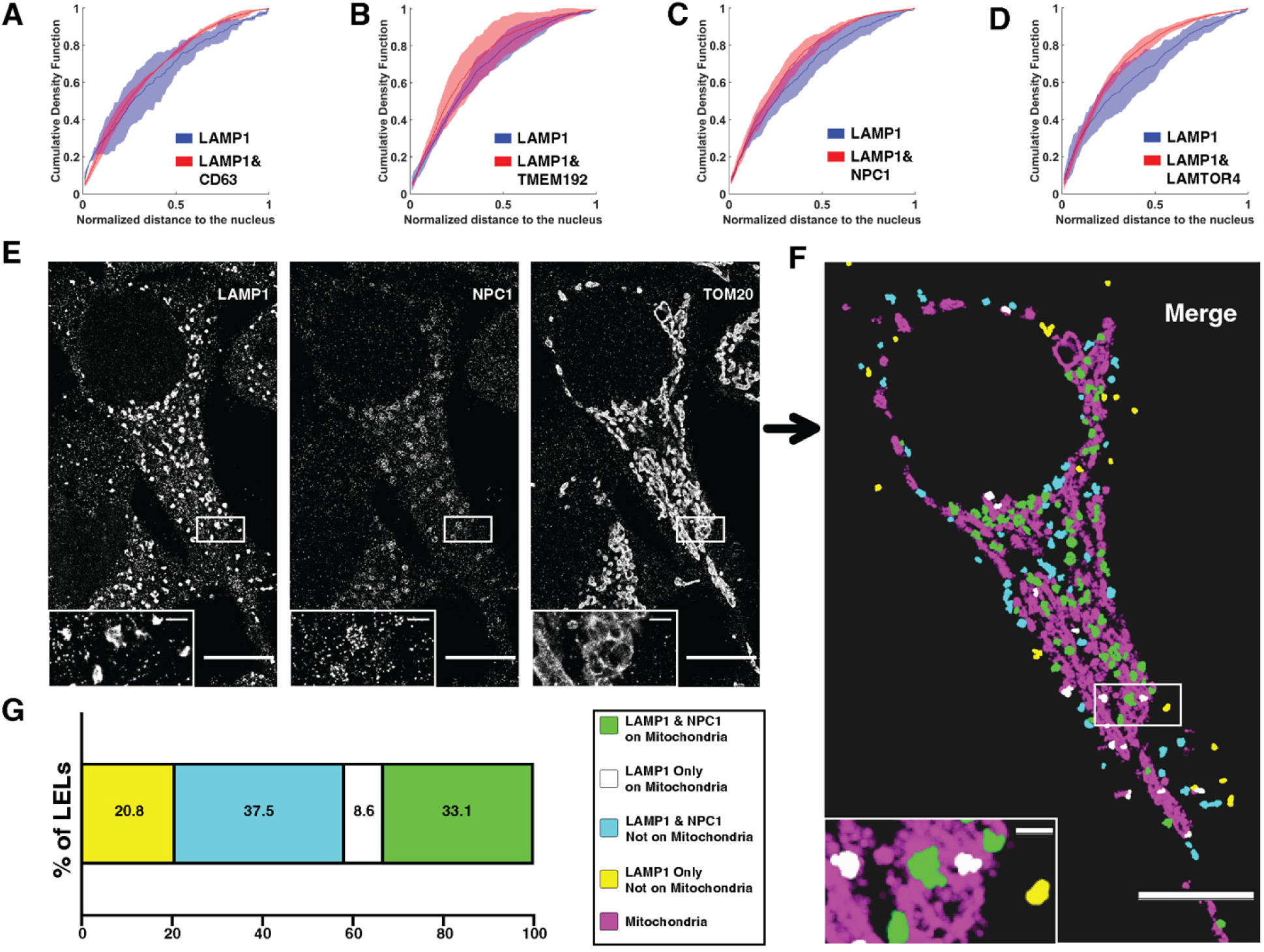
Spatial analysis reveals the location of LEL subsets with respect to the nucleus and mitochondria. **A-D:** Cumulative density function plots of LEL distance from the nucleus calculated from dual-color DNA-PAINT images in HeLa cells for subpopulations containing LAMP1 only or LAMP1 and target protein: CD63 **(A)**, TMEM192 **(B)**, NPC1 **(C)**, and LAMTOR4 **(D)**. Distance to the nucleus was normalized per cell to the maximum distance from the nucleus of an LEL in that cell, with values closest to zero indicating greatest proximity to the nucleus. Line indicates median with standard deviation between biological replicates. Kolmogorov-Smirnov tests were performed on the mean distributions from 3 independent biological replicates: for CD63 n = 14 cells, P = 0.261, no significance; for TMEM192 n = 13 cells, P = 0.794, no significance; for NPC1 n = 16 cells, P =0.261, no significance; for LAMTOR4 n = 16 cells, P = 0.069, no significance. **E-G:** 3-color DNA-PAINT image of LAMP1, NPC1, and mitochondria (TOM20) in HeLa cells. Raw images **(E)** and post-processed image **(F)** showing a spatial map of LELs with or without NPC1 in relation to mitochondria. Quantification **(G)** shows combined subpopulations of 5 cells from 3 biological replicates. Cell scale bars, 10 µm. Inset scale bars, 1 µm.

Considering NPC1’s crucial role in cholesterol export and the established proximity of lysosomes to other cellular organelles, we conducted 3-color multiplexed DNA-PAINT imaging to investigate the likelihood of NPC1-positive LELs being in close proximity to mitochondria (**Figure 4E, Supplementary Figure 5E**). We selected mitochondria for this analysis due to their known frequent interactions with lysosomes (Cisneros et al., 2022; Wong et al., 2018) and the availability of specific and high-affinity mitochondrial antibodies that effectively label mitochondrial membranes. We then adapted our co-localization analysis to differentiate between LELs with partial mitochondrial overlap and those showing no mitochondrial overlap (see Methods and **Figure 4E, F and Supplementary Figure 5E, F**). We note that partial overlap in our images does not imply the presence of LEL-mitochondria membrane contact, as the spatial resolution of our DNA-PAINT imaging is not sufficient to infer membrane contact sites. We spatially mapped and visualized the sub-populations within each individual cell, providing a depiction of both their unique protein composition and spatial relationship to mitochondria (**Figure 4F and Supplementary Figure 5F**). The analysis revealed that in HeLa cells approximately 50% of NPC1-positive LELs were overlapped with mitochondria, in contrast to only 30% of NPC1-negative LELs exhibiting mitochondrial overlap (**Figure 4G**). These findings suggest a higher propensity for NPC1-positive LELs to be in close spatial proximity of mitochondria in HeLa cells, although the functional relevance of this is not clear. Interestingly, this trend was not true in ARPE-19 cells (**Supplementary Figure 5G**), showing cell type specific sub-cellular positioning of distinct LEL subpopulations with respect to other organelles. These results further highlight the power of our quantitative pipeline in uncovering the intricate inter-relationships among various molecularly distinct organelle subpopulations.

### Higher-order multiplexing reveals molecularly distinct LEL subsets

Given that only specific subsets of LAMP1-positive LELs carry certain LEL proteins, we hypothesized that employing higher-order multiplexing with these markers could uncover the diversity within LEL subpopulations. However, one major hurdle in multiplexing beyond two or three targets is the limited availability of high-quality antibodies from unique species. Since most validated LEL antibodies are produced in rabbit, their combination in multiplexed immunofluorescence labeling and imaging presents a challenge. To address this, we leveraged a newly developed workflow where DNA-PAINT labeled anti-rabbit nanobodies, each tagged with unique oligo barcodes, are pre-incubated with their respective primary antibodies to create a stable antibody-nanobody complex (Sograte-Idrissi et al., 2020). We first validated that this protocol did not lead to target crosstalk by targeting two distinct structures – mitochondria and microtubules (**Supplementary Figure 5H**). We indeed obtained no discernable crosstalk between these two structures, validating that the antibody-nanobody complex is stable through the immunolabeling and imaging timeframe.

Another challenge associated with multiplexed imaging is the fact that only two spectrally distinct fluorophores are routinely used for DNA-PAINT, necessitating the removal and exchange of imager oligos between imaging sessions. To maintain precise alignment between sequentially imaged targets, we employed fluorescent beads as fiducial markers and conducted post-imaging alignment (**Supplementary Figure 5I**).

We next applied this approach to examine protein targets in HeLa cells that exhibited heterogeneous co-localization levels with LAMP1 (i.e. NPC1, and LAMTOR4) (**Figure 5A-C**). A similar approach was applied to ARPE-19 cells with three proteins that showed heterogeneous co-localization with LAMP1 (i.e. CD63, NPC1 and LAMTOR4) (**Figure 5D-F**). Our co-localization analysis once again enabled us to spatially map the unique LEL sub-populations within each individual cell (**Figure 5B, E**). In HeLa cells, the predominant LEL subpopulation comprised all three proteins (approximately 40% contained LAMP1, NPC1, and LAMTOR4) (**Figure 5C**). Given the 93.5±7.8% co-localization of LAMP2 with LAMP1 and 87±6.8% co-localization of CD63 with LAMP1 in HeLa cells, this subpopulation most likely also contains LAMP2 and CD63. However, we identified a significant (∼27%) subpopulation of LELs that were solely positive for LAMP1 (and presumably LAMP2/CD63) but lacked both NPC1 and LAMTOR4. These findings imply that LEL proteins typically identified in proteomic studies are not uniformly present on every LEL subpopulation, revealing significant heterogeneity in the protein composition of canonical LAMP1/2-positive LELs.

**Figure 5:**
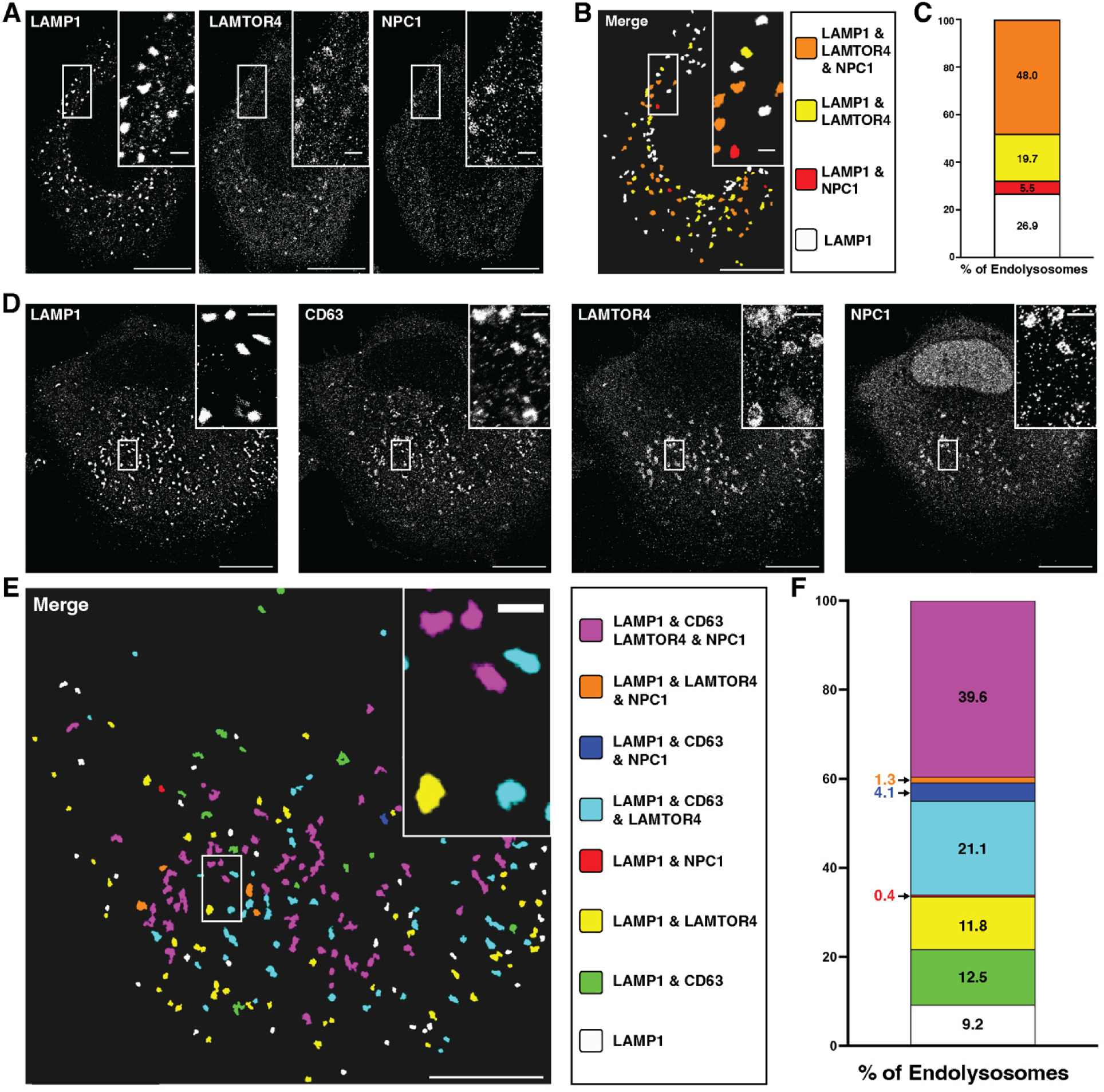
Higher-order multiplexing reveals molecularly distinct LEL subsets. **A-C:** 3-color DNA-PAINT image of LAMP1, LAMTOR4, and NPC1 in HeLa cells reveals four distinct subpopulations. Raw images **(A)** and post-processed image **(B)** showing a spatial map of these four subsets. Quantification **(C)** shows combined subpopulations of 4 cells from 3 biological replicates. Cell scale bars, 10 µm. Inset scale bars, 1 µm. **D-F:** 4-color DNA-PAINT image of LAMP1, CD63, LAMTOR4, and NPC1 in ARPE-19 cells reveals eight distinct subpopulations. Raw images **(D)** and post-processed image **(E)** showing a spatial map of these eight subsets. Quantification **(F)** shows combined subpopulations of 4 cells from 3 biological replicates. Cell scale bars, 10 µm. Inset scale bars, 1 µm.

This diversity was similarly observed in ARPE-19 cells, where cells could display up to eight distinct LEL subpopulations based on their protein makeup (**Figure 5E, F**). The most common subpopulation again included all four proteins (40% of LELs had LAMP1, CD63, LAMTOR, and NPC1) (**Figure 5F**). However, there were also subpopulations missing one to three of the examined proteins. Furthermore, we observed variability among individual cells in terms of these subpopulations, where certain cells were devoid of specific minority subpopulations, potentially indicating these subpopulations may lack significant functional importance or relevance.

## Discussion

In this study, we introduce a novel application of multiplexed and quantitative DNA-PAINT imaging as a robust and effective method for determining the heterogeneity of individual organelles in situ and under native conditions. While multiplexed DNA-PAINT has very recently been applied to visualize synaptic protein heterogeneity (Unterauer E. M. et al., 2023) as well as distribution of Golgi protein complexes (Schueder F. et al., 2023) to our knowledge, this is the first application of this approach to visualize the heterogeneity of native organelle sub-populations in the cellular context. Our work provides several key resources including: i) thoroughly validated antibodies that can be used for super-resolution visualization of high and low abundance LEL membrane proteins, ii) a robust, object-based co-localization analysis, enabling the determination of both the percentage of co-localization among organelles and the density of proteins on their membranes, and iii) comprehensive datasets that quantitatively profile six LEL membrane proteins across two different cell types.

Using this approach, we found that abundant and canonical LEL proteins (LAMP1 and LAMP2) exhibit a high degree of co-localization on the same compartments. Thus, these proteins act as general markers of the LEL population. Despite their prevalence across most LELs, we observed significant heterogeneity in the density of these proteins among individual LELs within a single cell. Furthermore, there was no correlation between the densities of different proteins, such as LAMP1 and LAMP2, suggesting that this heterogeneity might result from the stochastic integration of proteins onto the LEL membrane during their biogenesis or maturation. It would be interesting to explore in the future if this heterogeneity has any functional consequences. Beyond the variability in protein density, we also identified that LELs segregate into distinct sub-populations based on the presence or absence of other membrane proteins like TMEM192, NPC1, and LAMTOR4. In both HeLa and ARPE19 cells, we were able to identify various LEL sub-populations defined by their unique combinations of membrane proteins. Notably, the largest sub-populations were those containing all the visualized proteins (LAMP1/NPC1/LAMTOR4 in HeLa cells and LAMP1/CD63/NPC1/LAMTOR4 in ARPE-19 cells). This group, comprising a comprehensive array of canonical LEL proteins identified in proteomic studies, may represent a functionally distinct subset of LELs, potentially involved in NPC1-mediated cholesterol homeostasis and mTOR signaling. More work is needed to uncover the functional significance of these sub-populations and their distinct sub-cellular dynamics.

Employing correlative live-cell and super-resolution imaging (Balint et al., 2013; Verdeny-Vilanova et al., 2017) with sensors for autophagy (Kim and Seong, 2021), mTOR activity (Zhou et al., 2015), and luminal pH (Chin et al., 2021) could reveal more about these relationships. Previous studies in neurons showed that not all LAMP1-positive LELs contain degradative enzymes like cathepsin D. (Cheng et al., 2018). Additionally, recent work uncovered two distinct LELs characterized by their distinct sub-cellular positioning, morphology, lipid composition and mTORC1 activity (Ebner et al., 2023). These studies support the notion of functionally distinct LEL subsets. Our work further extends these studies by uncovering many additional LEL subpopulations based on their membrane composition. Recent proteomic studies revealed the heterogeneity of lysosome proteome between different tissues (Yu et al., 2024). In the future, it would be interesting to explore if specific LEL subpopulations are also differentially enriched in distinct tissues and cell types and if these subpopulations are differentially susceptible in aging and disease.

Proteins like NPC1 and LAMTOR4 exhibited distinct nanoscale organization patterns on the LEL membranes. NPC1 tended to form small, tightly packed nanodomains, while LAMTOR4 organized into larger, less dense nanodomains. Understanding the mechanisms that govern the organization of these proteins into these specific nanoscale platforms is crucial for determining whether their spatial organization influences their function. For example, NPC1, which plays a key role in cholesterol homeostasis, might segregate into cholesterol-enriched lipid domains (Wang et al., 2020a) on the LEL membrane. This spatial arrangement could be linked to lysosomal contact sites with other organelles including the ER, peroxisomes, Golgi and mitochondria to facilitate cholesterol delivery to these organelles (Radulovic et al., 2022). Additionally, LAMTOR4 is important for mTORC1 recruitment to the LEL membrane (Sancak et al., 2010) and the LAMTOR4 nanodomains may serve as platforms for efficient mTORC1 recruitment.

Our results also revealed that over-expression of LEL proteins can subtly yet significantly alter various characteristics of LELs, changes that might be challenging to detect with conventional, diffraction-limited microscopy. For example, over-expressing LAMP1 resulted in a small but significant enlargement of LELs, consequently reducing the surface density of LAMP2 and NPC1 proteins on their membranes. Additionally, LAMP1 over-expression influenced protein clustering, leading to the enlargement of NPC1 nanodomains with a reduced density of NPC1 proteins. These observations suggest the need for caution in interpreting data from over-expression experiments, as they can subtly but significantly affect lysosomal biology. In addition, we showed that proteins like TMEM192 and NPC1 are present only on a limited subset of LAMP1-positive LELs, potentially indicating unique properties and functions for these subpopulations. TMEM192 overexpression is used in Lyso-IP studies to biochemically isolate lysosomes for proteomic and metabolomic analysis. We showed that when over-expressed, this protein associates with all LAMP1-positive LELs, as previously reported (Abu-Remaileh et al., 2017). These results highlight a limitation of current Lyso-IP approaches, namely the inability to isolate and analyze distinct LEL sub-populations. Consequently, proteomic and metabolomic analyses fail to differentiate between molecularly diverse LEL sub-populations. To address this limitation, new strategies are required. One potential approach is the endogenous tagging of LEL proteins, which are specific to certain sub-populations, using CRISPR tagging techniques (Chen et al., 2018). This method could facilitate the isolation and comparative study of different LEL sub-populations, offering deeper insights into their distinct functions and properties.

Here, we profiled 6 LEL proteins, though proteomic studies have identified hundreds of LEL proteins including ion channels, transporters and components of the mTOR signaling pathway (Bagshaw et al., 2005; Chapel et al., 2013; Lubke et al., 2009; Schroder et al., 2010). The main limitation in expanding our multiplexing approach to a larger set of proteins is the scarcity of high-quality antibodies that specifically bind to these LEL proteins for endogenous level labeling. We evaluated a wide range of commercially available antibodies against numerous LEL proteins, many of which did not meet our stringent validation criteria due to lack of specificity. Therefore, to broaden our investigation to include more targets, there is a critical need for the development of new, high-quality LEL labeling reagents suitable for multiplexed DNA-PAINT. Emerging advancements in the creation of synthetic nanobodies (or sybodies) (Misson Mindrebo et al., 2023; Zimmermann et al., 2020) are particularly promising in this context. With the availability of such advanced reagents, our pipeline will be broadly applicable to profiling organelle heterogeneity at an unprecedented level of detail in the future.

## Methods

### Cell Culture

Wild type HeLa (ATCC CCL-2) and ARPE19 (ATCC CRL-2302) cell lines were obtained from the American Type Culture Collection (ATCC, Manassas, VA). The HeLa NPC1-null cell line was a kind gift from Prof. Neale Ridgway, Ph.D. (Dalhousie University, Halifax, Nova Scotia, Canada) (Zhao and Ridgway, 2017). HeLa cells were propagated in DMEM, and ARPE-19 cells in DMEM:F12 media. All culture media (GIBCO Laboratories, Grand Island, NY) was supplemented with 10% (vol/vol) fetal bovine serum and antibiotics. Cells were maintained at 37°C in the presence of 5% CO_2_ and routinely checked for mycoplasma contamination. To overexpress or knockdown a protein of interest, cells were transiently transfected at 50-60% confluency with plasmid expressing protein of interest (Table 1) and/or target specific siRNA using Lipofectamine 2000 reagent (Invitrogen) according to the manufacturer’s protocol. Cells were subjected to experimental treatments 24 h after transfection.

**Table 1:** Plasmids.

| Plasmid name | Catalog Number | Insert | Species |
| --- | --- | --- | --- |
| LAMP1-mGFP | Addgene# 34831 | LAMP1 | H. sapiens (human) |
| pRK5-LAMP1-FLAG | Addgene# 71868 | LAMP1 | H. sapiens (human) |
| LAMP2-BFP | Gift from Holzbaur lab (University of Pennsylvania) (Gallagher and Holzbaur, 2023) | LAMP2 | H. sapiens (human) |
| pMEH371 CD63 HA | Addgene# 168220 | CD63 | H. sapiens (human) |
| pLJC5-Tmem192-3xHA | Addgene# 102930 | TMEM192 | H. sapiens (human) |
| pRK5-FLAG-C7orf59 | Addgene# 42332 | LAMTOR4 | H. sapiens (human) |

### Western Blot Analysis

Western blot analysis was performed using the two-color Odyssey LI-COR (Lincoln, NE) technique according to the manufacturer’s protocol. All antibodies and corresponding dilutions used can be found in Table 2.

**Table 2:** Antibodies and DNA Imager Oligomers.

| Antibody/Oligo | Catalog Number | Experiments Used | Dilutions |
| --- | --- | --- | --- |
| Sheep Anti-LAMP1 | R&D Systems #AF4800 | DNA-PAINT; Widefield Imaging | 1:100; 1:500 |
| Mouse Anti-LAMP2 | Santa Cruz Biotech #sc18822 | DNA-PAINT; Widefield Imaging | 1:100; 1:500 |
| Mouse Anti-CD63 | Novus Bio #NBP2-42225 | DNA-PAINT; Widefield Imaging | 1:100; 1:500 |
| Rabbit Anti-TMEM192 | Abcam #ab185545 | DNA-PAINT; Widefield Imaging; Western Blot | 1:25; 1:500; 1:1000 |
| Rabbit Anti-NPC1 | Novus Bio #NB-400-148 | DNA-PAINT | 1:100 |
| Rabbit Anti-LAMTOR4 | Cell Signaling #13140 | DNA-PAINT; Widefield Imaging | 1:100; 1:500 |
| Mouse Anti-TOM20 | Santa Cruz Biotech #sc-17764 | DNA-PAINT | 1:100 |
| Mouse Anti-Flag | Millipore Sigma #F1804 | Widefield Imaging | 1:500 |
| Rabbit Anti-HA | Abcam #ab9110 | Widefield Imaging | 1:500 |
| Rabbit Anti-EEA1 | Cell Signaling #3288 | DNA-PAINT | 1:100 |
| Rabbit Anti-TMEM192 (alternative antibody) | Thermo Fisher #PA5-118314 | DNA-PAINT | 1:100 |
| Rabbit Anti-NPC1 | Abcam #ab134113 | Western Blot | 1:1000 |
| Mouse Anti-GAPDH | Proteintech #60004-1-Ig | Western Blot | 1:1000 |
| Rabbit Anti-TOM20 | Proteintech #11802-1-AP | DNA-PAINT | 1:100 |
| Rabbit Anti-Tubulin, Acetylated | Millipore Sigma #T6793 | DNA-PAINT | 1:100 |
| Anti-Mouse IgG+ Docking Site 1 | Massive Photonics, Massive-AB 2-Plex | DNA-PAINT | 1:100 |
| Anti-Rabbit IgG+ Docking Site 2 | Massive Photonics Massive-AB 2-Plex | DNA-PAINT | 1:100 |
| Donkey Anti-Sheep IgG+ Docking Site E2 | Jackson ImmunoResearch #713-005-147 | DNA-PAINT | 1:100 |
| Tag-Q Anti-GFP Single-Domain Antibody+ Docking Site 3 | Massive Photonics, Massive-TagQ Anti-GFP | DNA-PAINT | 1:100 |
| Anti-Rabbit Single-Domain Antibody+ Docking Site F1 | Massive Photonics, Custom Order | DNA-PAINT | 1:25 |
| Anti-Rabbit Single-Domain Antibody+ Docking Site F2 | Massive Photonics, Massive SDAB Fast One-Plex | DNA-PAINT | 1:25 |
| Donkey Anti-Mouse IgG+ Alexa Fluor 647 | Thermo Fisher #A-31571 | Widefield Imaging | 1:500 |
| Goat Anti-Rabbit IgG+ Cy3 | Thermo Fisher #A10520 | Widefield Imaging | 1:500 |
| IRDye800CW Donkey anti-Rabbit | LI-COR #926-32213 | Western Blot | 1:10,000 |
| IRDye680RD Donkey anti-Mouse | LI-COR #926-68072 | Western Blot | 1:10,000 |
| Imager Oligo for Docking Site 1+ Cy3B or ATTO655 | Massive Photonics, Massive-AB 2-Plex | DNA-PAINT | 500 pM |
| Imager Oligo for Docking Site 2+ Cy3B or ATTO655 | Massive Photonics, Massive-AB 2-Plex | DNA-PAINT | 500 pM |
| Imager Oligo for Docking Site 3+ Cy3B or ATTO655 | Massive Photonics, Massive-TagQ Anti-GFP | DNA-PAINT | 100-300 pM |
| Imager Oligo for Docking Site E2+ Cy3 | IDT, Custom Order | DNA-PAINT | 500 pM |
| Imager Oligo for Docking Site F1+ Cy3B or ATTO655 | Massive Photonics, Custom Order | DNA-PAINT | 1.5 nM |
| Imager Oligo for Docking Site F2+ Cy3B or ATTO655 | Massive Photonics, Massive SDAB Fast One-Plex | DNA-PAINT | 1.5 nM |

### Sample Preparation

#### Generation of anti-sheep DNA-PAINT secondary antibodies

Affinipure donkey anti-sheep secondary antibody (Jackson Immunoresearch Labs) was conjugated to 5’-TTATCTACATA-3’ for DNA-PAINT. This docking site is referred to as docking site E2 and imaged with corresponding imager strand E2 (custom ordered from IDT, Coralville, Iowa). DNA was conjugated to the antibody via DBCO-sulfo-NHS ester chemistry according to the protocol described previously (Schnitzbauer et al., 2017). Briefly, the antibody was incubated with 10-fold excess of bifunctional DBCO-sulfo-NHS ester (Jena Bioscience, Cat.# CLK-A124-10) for 2 hours at +40C. Unbound linker was removed using Zeba Spin Desalting columns (0.5 mL, 7K MWCO; Thermo Fisher Scientific, 89882). Azide-modified DNA was added to the DBCO-antibody in a 15 molar excess and incubated for 1 hour at room temperature, protected from light. At the end of incubation, buffer was exchanged to PBS using Amicon centrifugal filters (100,000 MWCO). Antibody labeling was confirmed using the NanoDrop spectrophotometer by the shift of the peak signal from 280 nm towards 260 nm.

### Standard DNA-PAINT immunostaining

Cells were seeded onto LabTek-II imaging chambers (Nalge Nunc, Thermo Fisher) before fixation. Glyoxal was tested as a fixative for 10 minutes at room temperature (Richter et al., 2018) but reduced staining quality when compared to 4% paraformaldehyde (PFA) in PBS pre-warmed to 37°C for 20 minutes at room temperature, thus PFA was used for all experiments. Cells were then washed 3X with PBS and permeabilized for 10 minutes in 0.1% saponin in PBS. 0.2% Triton X-100 was tested as an alternative permeabilization, but reduced retention of target proteins on LEL membranes, thus saponin was used for all experiments. Blocking was done in Primary Blocking Buffer (10% donkey serum, 0.1% saponin, 0.05 mg/mL sonicated salmon sperm DNA (Stratagene, La Jolla, California) in PBS for 1 hour at room temperature. Following blocking, cells were incubated for one hour at room temperature in primary antibodies described in Table 2 diluted in Primary Blocking Buffer. After primary antibody staining, cells were washed briefly with 1X with PBS and 3X for 5 minutes with 1X Wash Buffer (Massive Photonics, Gräfelfing Germany). Secondary antibodies were diluted in Antibody Incubation Buffer (Massive Photonics) as described in Table 2 and incubated for 1 hour at room temperature. Subsequently, cells were washed briefly 1X with PBS, 3X for 7 minutes with 1X Wash Buffer (Massive Photonics), and 1X for 5 minutes with PBS and stored at 4 degrees in fresh PBS until imaging. For LAMP1 and LAMTOR4, staining quality was improved by using this protocol sequentially with a 15-minute post-fixation using 4% PFA between staining for each target. For LAMP1-mGFP overexpression samples, anti-GFP single-domain antibody was used to detect this population and was added during the secondary antibody incubation.

### Immunostaining for widefield imaging

Immunostaining for widefield imaging was done as above but with lower antibody concentrations (see Table 2) and slightly altered blocking buffers. It was not necessary to add sonicated salmon sperm DNA to blocking buffer to block off-target DNA-PAINT imager oligo binding in the nucleus, and thus primary and secondary antibody incubation was performed with antibodies diluted in Widefield Imaging Buffer (10% donkey serum, 0.1% saponin in PBS).

### DNA-PAINT immunostaining with nanobody pre-mixing

Multiplexed imaging of LAMTOR4 and NPC1 required an alternative staining protocol as the antibodies to both these targets were raised in rabbit. We utilized a previously described protocol (Sograte-Idrissi et al., 2020) for pre-mixing primary antibodies with secondary nanobodies to circumvent this issue. Excess DNA conjugated nanobodies at four-fold higher concentration than saturating levels of secondary antibody were pre-mixed with each primary antibody for 30 minutes at room temperature in PBS, before they were subsequently mixed and incubated with the cells during the secondary antibody step. To minimize the possibility of crosstalk as much as possible, these samples were imaged immediately following staining completion. As an additional validation, this protocol was tested on acetylated tubulin and TOM20 using rabbit primary antibodies for each (Table 2).

### Widefield imaging

Widefield images were acquired using a Nanoimager (ONI, Oxford, UK) equipped with 405-nm, 488-nm, 561-nm and 640-nm lasers, 498-551- and 576-620-nm band-pass filters in channel 1, 666-705-nm band-pass filters in channel 2, a 100X 1.45 NA oil immersion objective (Olympus), and a Hamamatsu Flash 4 V3 sCMOS camera. Widefield images were captured at 200 ms exposure using epifluorescence illumination and exported from the NimOS software (ONI) for visualization in FIJI (Schindelin et al., 2012). For visualization purposes only, a rolling ball background subtraction with a radius of 5 pixels was performed on these images.

### DNA-PAINT imaging

DNA-PAINT images were acquired on the same Nanoimager system as was used for widefield imaging, and all experiments were conducted at 30°C using HiLo illumination.

### Dual-color DNA-PAINT of LEL targets

Before imaging, appropriate imager oligo strands (see Table 2) for both labeled targets were diluted to a final concentration of 500 pM in Imaging Buffer (Massive Photonics) and added to imaging chambers. For imaging of LAMP1-GFP with GFP nanobody, 100-300 pM imager was used, as 500 pM yielded overlapping fluorescent signal and was deemed too high. For dual-color experiments, one imager with conjugated Cy3 or Cy3B and one imager with conjugated ATTO655 were used. The sample was imaged at 100 ms exposure using a laser program alternating between 561-nm excitation and 640-nm excitation every 100 frames for a total of 50,000 frames, 25,000 frames for each target.

### Three-color DNA-PAINT of LAMP1, NPC1, and TOM20

Before imaging, samples were incubated with 0.1 µm Tetraspeck microspheres (Invitrogen, Thermo Fisher) in PBS to use as a fiducial marker. Appropriate imager oligo strands for LAMP1 and NPC1 were diluted to 500 pM in Imaging Buffer and added to imaging chambers. The sample was imaged at 100 ms exposure using a laser program designed for sequential imaging of targets, beginning with 25,000 frames with 561-nm excitation and followed by 25,000 frames with 640-nm excitation. The third step in the laser program was 2,500-5,000 frames with no laser excitation to allow for imager oligo exchange for imaging of TOM20. The imager oligo solution on the sample was removed and the sample was washed 3X with PBS and replaced with 500 pM of the appropriate imager oligo strand for TOM20 conjugated to either Cy3B or ATTO655. The fourth step in the laser program turned on the appropriate laser corresponding to the fluorophore used (561-nm for Cy3B or 640-nm for ATTO655) for 10,000 frames.

### Higher-order multiplexed DNA-PAINT using secondary nanobodies

Before imaging, samples were incubated with 0.1 µm Tetraspeck microspheres in PBS to use as a fiducial marker. For both three-color imaging of LAMP1, NPC1, and LAMTOR4 in HeLa cells and four-color imaging of LAMP1, CD63, NPC1, and LAMTOR4 in ARPE-19 cells, appropriate imager oligos for LAMTOR4 and NPC1 were first diluted to 500 pM in Imaging Buffer and added to the imaging chambers. The sample was imaged at 50 ms exposure, per manufacturer recommendation for the nanobodies, with a laser program designed for sequential imaging of targets. First, LAMTOR4 was imaged for 25,000 frames. Next, NPC1 was imaged for 50,000 frames. The third step in the laser program was 5,000 frames with no lasers enabled to allow for imager oligo exchange to image remaining targets. The imager oligo sample on the stage was removed and the sample was washed 3X with PBS and replaced with 500 pM of appropriate imager oligo for LAMP1 in HeLa cells, or LAMP1 and CD63 in ARPE-19 cells. The fourth step in the laser program imaged LAMP1 for 25,000 frames, after which three-color imaging in HeLa cells was completed. For four color imaging in ARPE-19 cells, a fifth step imaged CD63 for 25,000 frames.

### DNA-PAINT image analysis

For all experiments, data were exported from NimOS software (ONI) with the following initial filtering parameters applied: Photon counts above 300, an X/Y localization precision of 30 nm, and a Sigma X/Y between 0-250 nm. These parameters were identical for all targets in all experiments. All following analyses were performed in MATLAB R2022b (The MathWorks, USA) using custom-made code (Github: https://github.com/LakGroup/Data_Analysis_Browser_Bond_etal).

### Segmentation and clustering

Individual cells were cropped, and reference channel localizations were Voronoi segmented and clustered with a minimum of 25 localizations and a maximum Voronoi area threshold, based on on which target protein was used as the reference channel and visual inspection of the segmentation quality. For LAMP1, thresholds ranged from 68-548 nm^2^. For LAMP2, thresholds ranged from 178-342 nm^2^. For LAMP1-GFP imaged with GFP-nanobody, thresholds ranged from 178-1,369 nm^2^. To remove background antibody signal, clusters were then filtered above a minimum clustered area of 49,912 nm^2^, which corresponds to a circle with a radius of 125 nm, as previous super-resolution imaging showed lysosome size ranges between 200-800 nm (Verdeny-Vilanova et al., 2017). For dual-color experiments with wildtype HeLa and ARPE-19 cells using all standard imaging parameters (100 ms exposure, 25,000 frames, 500 pM imager oligo) and LAMP1 as the reference, clusters were additionally filtered for a minimum of 2,500 localizations.

For experiments examining colocalization of LELs with mitochondria, TOM20 was used as a mitochondrial reference channel and Voronoi segmented and clustered with a threshold of 3,422 nm^2^ and post-processed for a minimum of 500 localizations and minimum clustered area of 49,912 nm^2^. The higher Voronoi threshold of 3,422 nm^2^ compared the one chosen for LAMP1/2 ensured that the mitochondria were clustered as the large network they represent in cells rather than separate segments.

### Estimating imaging completeness

The imaging of an organelle is considered complete if the area of the localizations that describe the organelle no longer changes with an increasing number of frames. This area evolution is considered in a pair-wise way, for each cluster of the reference channel (Ch_REF_) and the corresponding target channel (Ch_TARGET_) cluster.

In practice, the localizations in Ch_REF_ were Voronoi-segmented and clustered using the most broadly applicable parameters used in the colocalization analysis (LAMP1 clustering threshold of 178 nm^2^). The localizations in Ch_TARGET_ were not Voronoi-segmented and clustered, as segmentation of protein targets that show low density and sparse localizations is challenging. Instead, the localizations in this channel that were spatially located within a cluster of Ch_REF_ were considered as a single ‘cluster’. The evolution of the area of each individual cluster in Ch_REF_ (and its associated ‘cluster’ in Ch_TARGET_) was followed with respect to the number of frames (by considering the area of their alphashape object), beginning at the frame that had the first detected localization. Each curve was normalized to its final area and visualized in percentage, to remove size dependencies. For the localizations of the Ch_TARGET_ channel, the area evolution was visualized in terms of the final area of the associated organelle of Ch_REF_.

### Colocalization of LEL targets

To perform the colocalization analysis, the data in the reference channel (LAMP1 unless otherwise indicated; Ch_REF_) were considered as Voronoi-segmented and clustered data, whereas the data from the target channel (Ch_TARGET_) were considered as localizations, for the same reasons as described above.

First, the localizations in Ch_TARGET_ that are spatially located within the boundaries of Ch_REF_ clusters are grouped together and considered as a ‘cluster’. Then, the local background density distribution is calculated by considering the area that is not directly adjacent to the Ch_REF_ clusters, but proximal to them (i.e. between 234 and 585 nm around the border). The area adjacent to Ch_REF_ clusters (i.e., between 0-234 nm around the border) was not considered as some targets have a locally increased density in this region (i.e. LAMTOR4, discussed below). Ch_TARGET_ ‘clusters’ are then considered to be statistically significant clusters and colocalized if they have a density that is larger than 3 standard deviations higher than the mean of the local background distribution.

Additional output of this analysis is given as properties of the clusters. For the individual clusters in Ch_REF_, the area, the number of localizations, its density, and its distance to the nucleus (see next section) are calculated. For the colocalized clusters of Ch_TARGET_, the number of localizations and its density are calculated.

For all experiments using LAMTOR4 as Ch_TARGET_, a small alteration was implemented, as LAMTOR4 is part of a protein complex that sits on the lysosomal membrane rather than an integral membrane protein, and thus the localization pattern from this target showed a clear bias to extend beyond the edges of the LAMP1 Ch_REF_ cluster. To accommodate for this, the clusters of Ch_REF_ were slightly expanded (with 20 nm), but the rest of the analysis described above remained the same.

### Distance from nucleus analysis

The location of the nuclei could easily be discerned visually from the DNA-PAINT images and were therefore manually annotated. Their coordinates were then extracted and used to determine the distance of the clusters in Ch_REF_ to this nucleus (nearest border to border distance). In rare occurrences where an LEL was inside the nucleus area, its distance was considered as 0.

To accommodate for different cell sizes, the distances of the individual lysosomes to the nucleus were normalized between 0 and 1 by dividing by the maximum distance in each cell. This was appropriate as all cells showed a distribution of LEL spread throughout the entire cell spanning both the perinuclear and peripheral regions. Then, the data were split up into the two colocalized categories (see previous section; colocalized vs non-colocalized clusters), and a histogram of distances for each category was constructed using 10% bins by pooling data from all cells.

### Colocalization of LELs with mitochondria

For analysis of LEL overlap with mitochondria, it is possible to segment and cluster the data from both mitochondrial (TOM20) and LEL reference (LAMP1) channels, and a different strategy than previously described can be used for the colocalization analysis. Here, all channels have clearly clustered localizations, and thus a polyshape/alphashape can be constructed for the clusters in both the reference channel (TOM20, Ch_REF_) and the target channel (LAMP1, Ch_TARGET_). These objects can then be compared to each other to determine the degree of overlap between them and are considered as colocalized when there is at least 10% overlap between them. The percentage of overlap is defined as the percentage of localizations of the Ch_TARGET_ cluster that is spatially located within a cluster of the Ch_REF_ channel with respect to its total number of localizations. In the case that a Ch_TARGET_ cluster overlaps with multiple Ch_REF_ clusters, it is associated to the Ch_REF_ cluster that it overlaps the most with.

### Drift correction for higher-order multiplexing

For 3- and 4-color DNA-PAINT acquisitions, drift correction was performed after export from the NimOS software to ensure each channel was drift corrected separately and any changes in localization not due to drift (i.e. small changes in position during manual imager swaps) were not improperly corrected (see below section on alignment). To correct for drift throughout the measurement, the Drift at Minimum Entropy (DME) method was used (Cnossen et al., 2021). In brief, drift will increase the reconstruction uncertainty in localization microscopy, and therefore the entropy will increase as well. This property is associated to the fact that the localization microscopy reconstruction is an approximation of the true distribution of the molecules (according to a probability distribution). Thus, to remove the drift from the localized molecules, the entropy of this reconstruction can be minimized, which can be estimated by minimizing an upper bound of the statistical entropy of the probability distribution of all the localizations. For more details about practical implementations, we refer to the original work (Cnossen et al., 2021). One small change was made to the original algorithm which was to calculate the drift with respect to the first frame for each channel, rather than with respect to the mean drift.

### Channel alignment for higher-order multiplexing

For 3- and 4-color measurements, a change of imager had to be made in the middle of the measurement. Due to the manual intervention, a slight misalignment of the sample (before vs after) can occur that could not be corrected using the drift correction method described above as the change was abrupt and not gradual. To correct for this, Tetraspeck microspheres were included during sample preparation that emit fluorescent signal in each channel that was measured for the entire acquisition duration. The localizations of this signal could then be used to align the channels correctly and remove the disruption introduced by the manual intervention. To align the channels, beads were manually selected and the distance between the centroids of the beads (in a pairwise way) was then calculated. These distances were then averaged over the selected beads and the channels were corrected accordingly.

### Pattern analysis of NPC1 and LAMTOR4 on LEL membranes

To analyze the clustering of proteins on LEL membranes, we used DBSCAN clustering on the co-localized localizations of Ch_TARGET_ (Ester et al., 1996), given a search radius and a minimum number of points. The appropriate search radius was determined for each target using visual inspection as follows: NPC1 in wild-type HeLa cells 17.6 nm, NPC1 in LAMP1 overexpressing HeLa cells 23.4 nm, and LAMTOR4 in wild-type HeLa cells 23.4 nm, and a minimum number of points of 5. Clusters were post-filtered for a minimum of 10 localizations per cluster for all targets.

Once the localizations were clustered, we additionally determined the number of localizations per cluster, cluster area, cluster density and cluster diameter.

## Acknowledgements

We would like to thank Qing Tang, Patricia Colosi, Elizabeth Gallagher, Vera Moiseenkova-Bell and Claire H. Mitchell at the University of Pennsylvania for critical reading and detailed comments on the manuscript. This work is funded by R01 GM133842 (to M.L.), RM1 GM136511 (to M.L.), CMMI-1548571 (to M.L.).

## Data Availability Statement

The raw imaging data is available upon request as it is too large to deposit in a data repository. All other data used in making the plots in the main and supplementary figures have been deposited to Figshare (10.6084/m9.figshare.25429189).

## Code Availability Statement

The custom written Matlab analysis code has been deposited on Github (https://github.com/LakGroup/Data_Analysis_Browser_Bond_etal).

## Author Contributions

C.B. and J.X acquired and analyzed data. E.M.S. carried out transfection and western blot experiments and provided reagents including custom-labeled antibodies. S.H. developed custom written data analysis code. M.L. supervised the work and acquired funding. M.L. and C.B. wrote the manuscript. All authors provided feedback on the manuscript.

