## Supplementary Information for "Multiplexed DNA-PAINT Imaging of the Heterogeneity of Late Endosome/Lysosome Protein Composition"

**Supplementary Figures:**

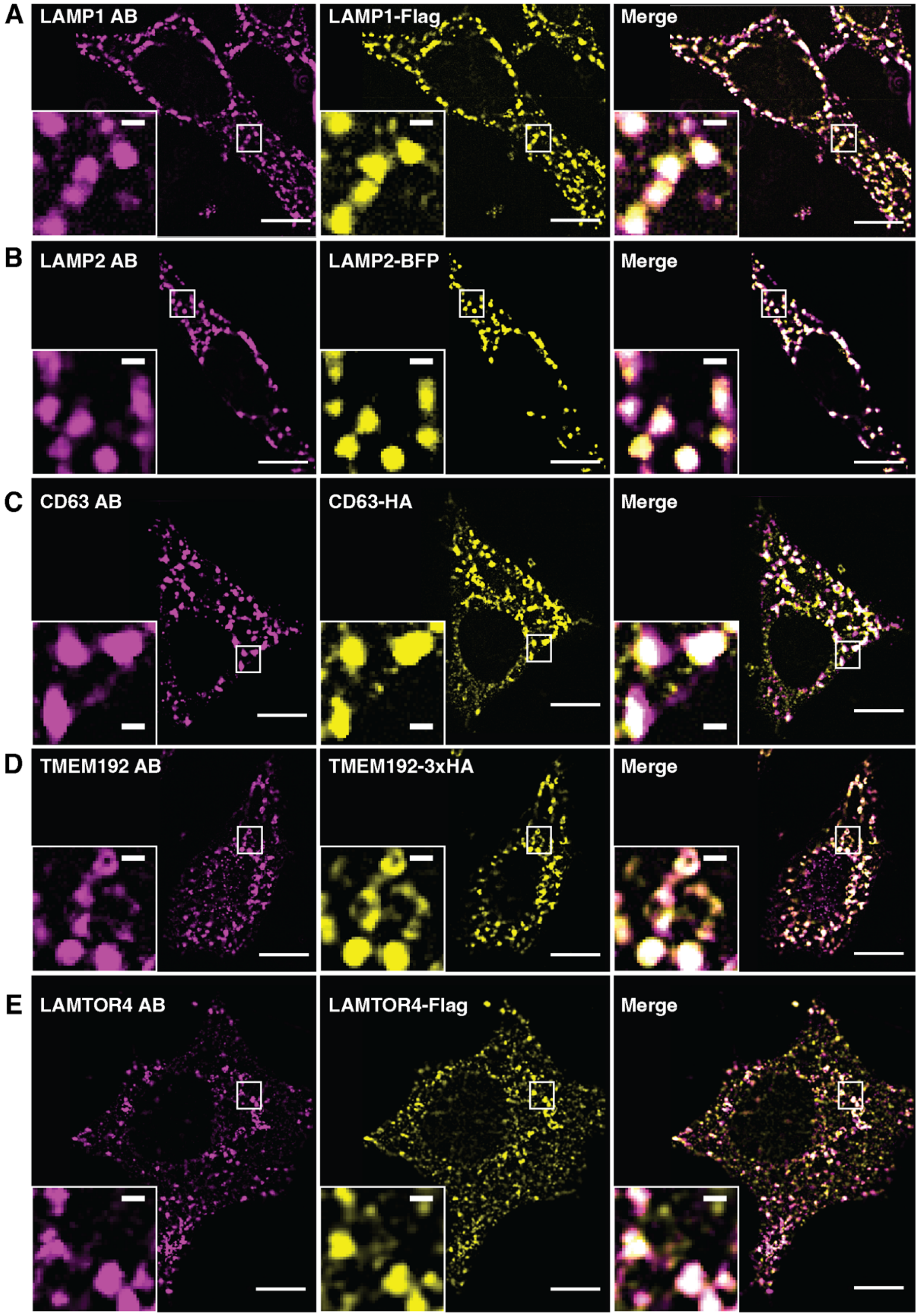


**Supplementary Figure 1: Overexpression of target proteins confirms antibody specificity.** Overexpression of tagged constructs of **A.** LAMP1-Flag **B.** LAMP2-BFP **C.** CD63-HA **D.** TMEM192-3xHA and **E.** LAMTOR4-Flag were used to verify antibody specificity in HeLa cells. Widefield images showing the co-staining with antibodies to the target protein (magenta) and the overexpression tag (yellow) colocalized for all proteins listed. Cell scale bars, 10 µm. Inset scale bars, 1 µm.

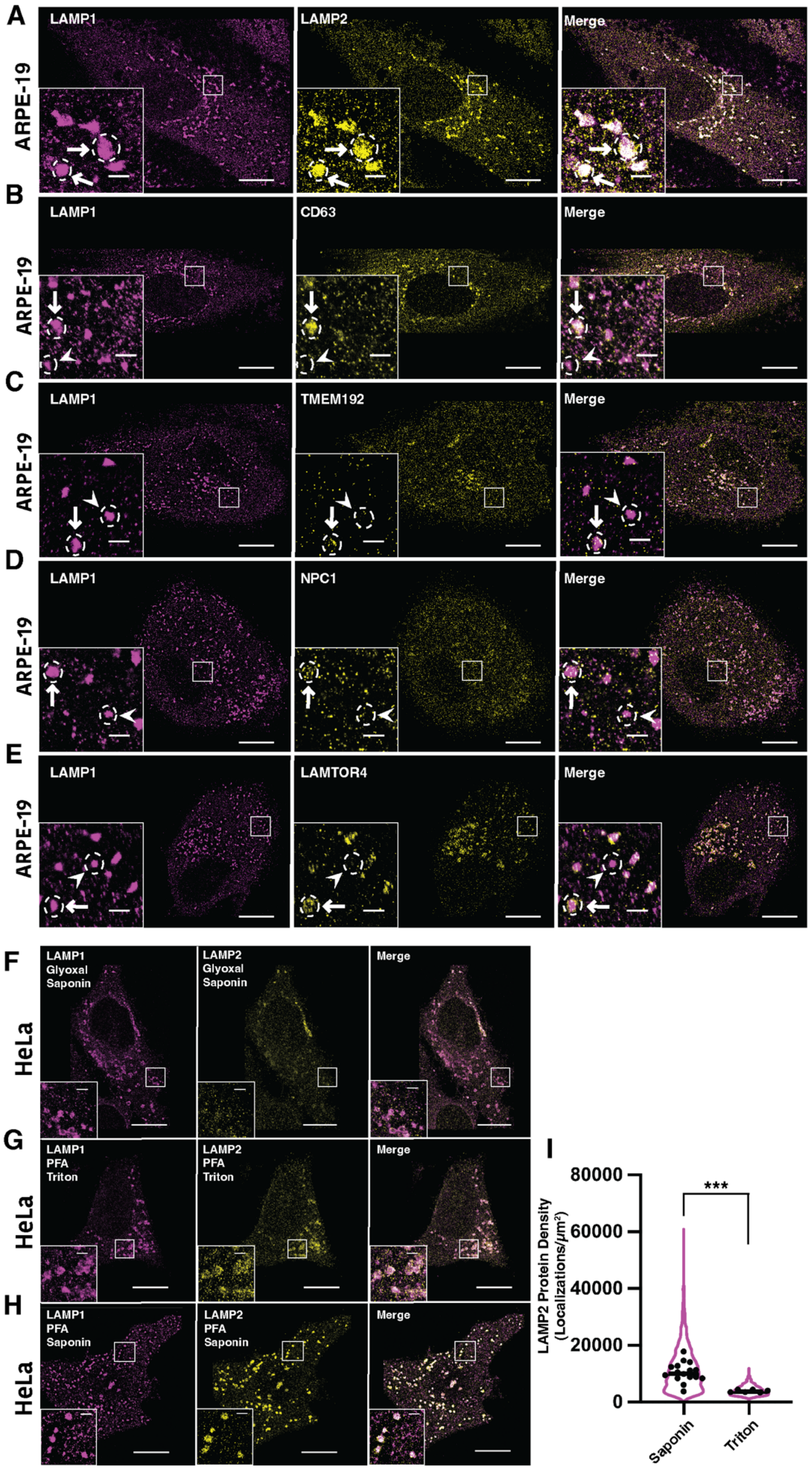

A
LAMP1
LAMP2
Merge
ARPE-19
B
LAMP1
CD63
Merge
ARPE-19
C
LAMP1
TMEM192
Merge
ARPE-19
D
LAMP1
NPC1
Merge
ARPE-19
E
LAMP1
LAMTOR4
Merge
ARPE-19
F
LAMP1 Glyoxal Saponin
LAMP2 Glyoxal Saponin
Merge
HeLa
G
LAMP1 PFA Triton
LAMP2 PFA Triton
Merge
HeLa
H
LAMP1 PFA Saponin
LAMP2 PFA Saponin
Merge
HeLa
I
LAMP2 Protein Density (Localizations/μm²)
80000
60000
40000
20000
0
***
Saponin
Triton

**Supplementary Figure 2: Dual-color DNA-PAINT reveals LEL subpopulations in ARPE-19 cells, as well as optimal fixation and permeabilization conditions.**

**A-E:** Representative DNA-PAINT images of LAMP1 reference channel (magenta) and target protein channel (yellow) in ARPE-19 cells for **A.** LAMP2 **B.** CD63 **C.** TMEM192 **D.** NPC1 and **E.** LAMTOR4. Arrows indicate LELs positive for both LAMP1 and target protein, arrowheads indicate LELs positive for LAMP1 and negative for target protein, dotted circle indicates which LEL arrow or arrowhead refers to. Cell scale bars, 10 µm. Inset scale bars, 1 µm.

**F:** Representative DNA-PAINT image of glyoxal-fixed HeLa cells stained for LAMP1 and LAMP2 shows significant disruption of LAMP2 staining.

**G-I:** Representative DNA-PAINT image of HeLa cells fixed with warm 4% PFA and permeabilized with 0.2% Triton X-100 **(G)** or 0.1% saponin **(H)**. Quantitative analysis of protein density **(I)** shows a significant reduction with 0.2% Triton X-100. Mann Whitney U-Test was performed to compare LAMP2 protein density in saponin treated cells (N = 3 biological replicates, n = 16 cells) with Triton X-100 treated cells (N = 1 biological replicate, n = 6 cells) (P = 0.0002).

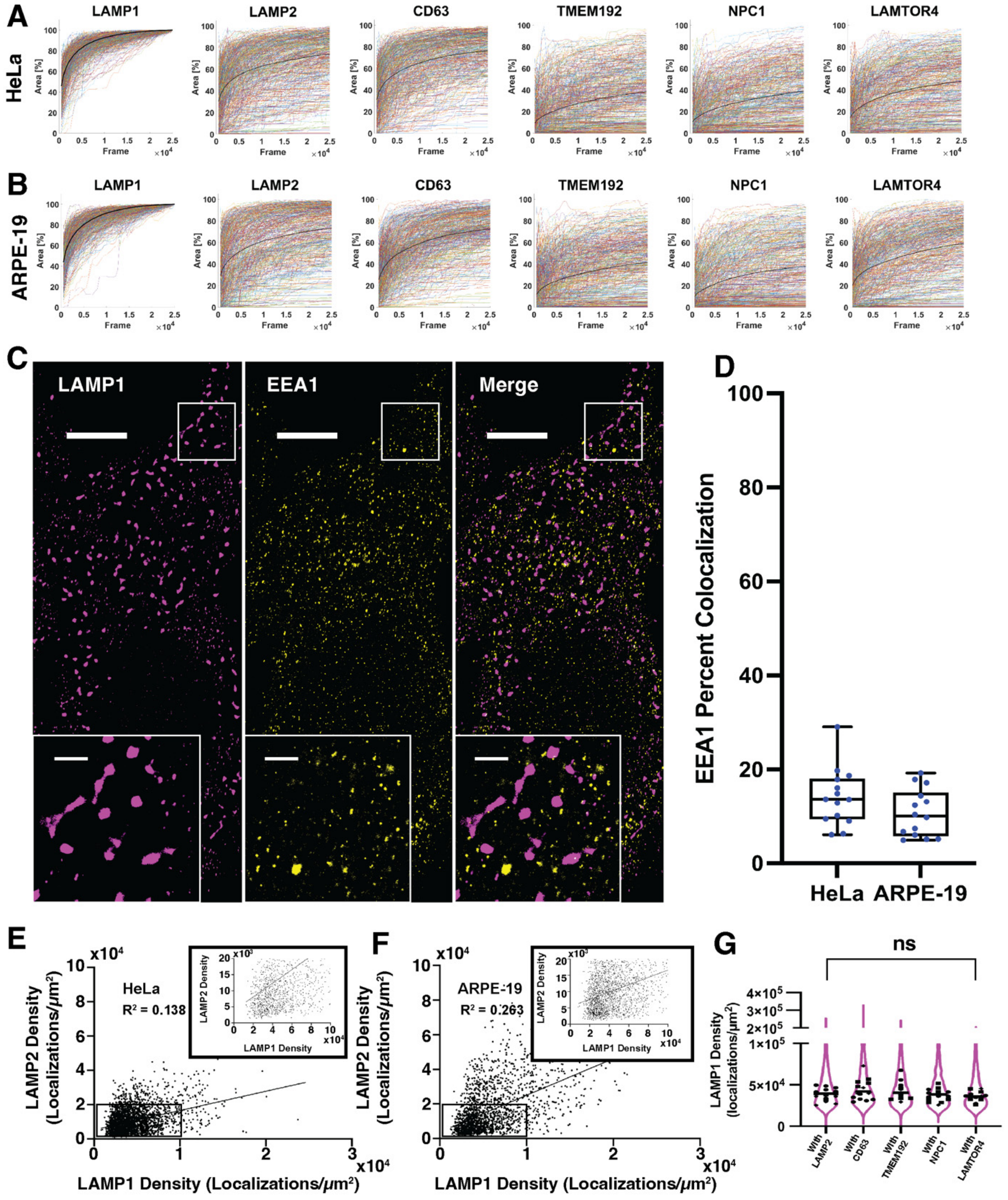


**Supplementary Figure 3: Controls demonstrate the robustness of DNA-PAINT imaging.**

**A-B:** Plots show the percent area of an LEL calculated from the reference image (LAMP1) covered by the localizations from the target protein in **A.** HeLa and **B.** ARPE-19 cells. Colored lines show traces from 500 randomly selected LELs per target. Solid black line indicates average over all traces for that target. As

LAMP1 was used as the reference channel, these traces always reach 100% coverage. Other targets cover a variable percent of LEL area, and importantly individual traces approach a plateau by 25,000 frames for all targets, indicating majority of target protein localizations have been captured by this imaging time.

**C-D:** Representative dual-color DNA-PAINT images and quantification of LAMP1 and EEA1 in HeLa cells shows distinct organelle populations labeled by each marker, indicating that LAMP1 does not label EEA1-positive early endosomes. N = 3 biological replicates per cell type, n = 14 HeLa cells, n = 14 ARPE-19 cells. Cell scale bars, 10 µm. Inset scale bars, 1 µm.

**E-F:** LAMP1 and LAMP2 levels on individual LELs in **E.** HeLa and **F.** ARPE-19 cells show minimal correlation. Pearson’s correlation coefficients ($R^2$) were 0.138 and 0.263 for HeLa and ARPE-19 cells, respectively.

**G:** LAMP1 localization density is consistent across multiple distinct dual-color experiments in HeLa cells. Kruskal-Wallis test was performed on median LAMP1 density per cell across experiments, P = 0.5342, not significant.

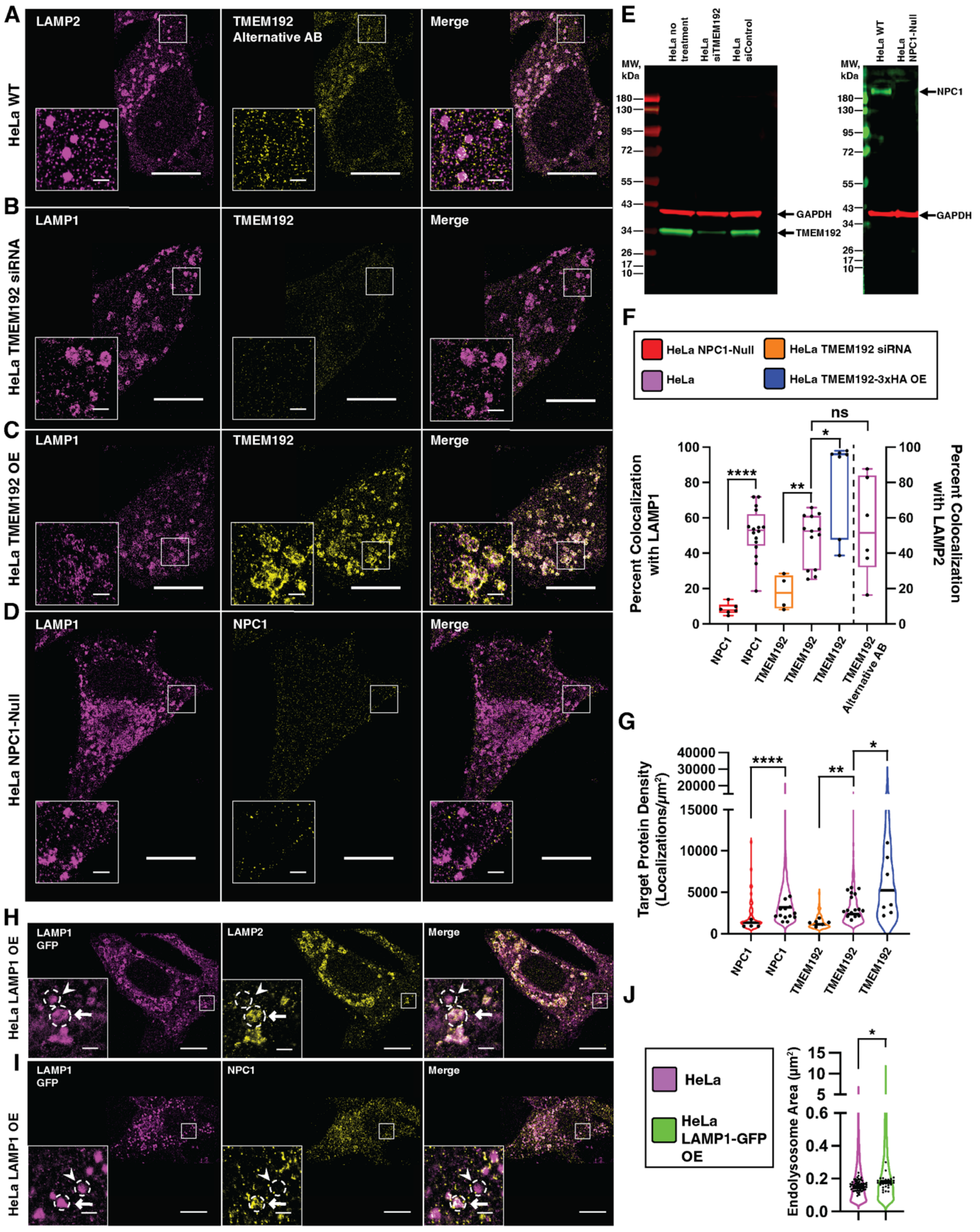

**Supplementary Figure 4: TMEM192 and NPC1 antibodies are robust and specific, and LAMP1-GFP overexpression in HeLa cells induces changes in LELs.**

**A:** Representative DNA-PAINT image of LAMP2 and TMEM192 in HeLa cells imaged using an alternative TMEM192 antibody shows a staining pattern consistent with results from the main antibody used. Cell scale bars, 10 µm. Inset scale bars, 1 µm.

**B:** Representative DNA-PAINT image of TMEM192 when the TMEM192 protein is knocked down via siRNA in HeLa cells shows a significant reduction in TMEM192 signal on LAMP1 with respect to wildtype conditions.

**C:** Representative DNA-PAINT image of TMEM192 in TMEM192-3xHA overexpressing HeLa cells shows near-complete overlap with LAMP1.

**D:** Representative DNA-PAINT image of NPC1 in NPC1-null HeLa cells shows minimal overall signal and overlap with LAMP1 with respect to wildtype conditions.

**E:** Western blot analyses of NPC1 and TMEM192 levels in HeLa cells. NPC1-null HeLa cells show loss of NPC1 signal. TMEM192 siRNA knockdown HeLa cells show a reduction in TMEM192 signal.

**F:** Quantification of co-localization of NPC1 and TMEM192 in HeLa cells. Plot line color indicates cell type, black circles indicate individual cells. Mann-Whitney U-tests were performed to compare: NPC1-null HeLa cells (N = 1 biological replicate, n=6 cells) with HeLa cells (N = 3 biological replicates, n=16 cells) (P < 0.0001), TMEM192 siRNA knockdown HeLa cells (N = 1 biological replicate, n=4 cells) with HeLa cells (N = 3 biological replicates, n=13 cells) (P = 0.0034 ), TMEM192-3xHA overexpressing cells (N = 1 biological replicate, n=7 cells) with wild type HeLa cells (N = 3 biological replicates, n=13 cells) (P = 0.0297), and wild type HeLa cells imaged with TMEM192 main antibody (N = 3 biological replicates, n=13 cells) or alternative antibody (N = 1 biological replicate, n=6 cells) (P = 0.6388, no significance).

**G:** Protein density of NPC1 and TMEM192 in HeLa cells. Plot line color indicates cell type, black line indicates median target density on LELs positive for a given target. Black circles represent individual cell medians. Mann-Whitney U-tests were performed to compare: NPC1-null HeLa cells (N = 1 biological replicate, n=6 cells) with wild type HeLa cells (N = 3 biological replicates, n=16 cells) (P < 0.0001), TMEM192 siRNA knockdown HeLa cells (N = 1 biological replicate, n=4 cells) with wild type HeLa cells (N = 3 biological replicates, n=13 cells) (P = 0.0034), and TMEM192 overexpressing cells (N = 1 biological replicate, n=7 cells) with wild type HeLa cells (N = 3 biological replicates, n=13 cells) (P = 0.0236).

**H-I:** Representative DNA-PAINT images of **H.** LAMP2 and **I.** NPC1 with LAMP1 in HeLa cells overexpressing LAMP1-GFP. Arrows indicate LELs positive for both LAMP1-GFP and target protein, arrowheads indicate LELs positive for LAMP1 and negative for target protein, dotted line indicates which LEL arrow or arrowhead refers to. Cell scale bars, 10 µm. Inset scale bars, 1 µm.

**J:** Comparison of LEL area in wild type vs. LAMP1-GFP overexpressing HeLa cells. Black circles indicate individual cell medians. Mann-Whitney U-test was performed to compare cell medians of HeLa cells (n=75) with LAMP1-GFP overexpressing HeLa cells (n=26) (P = 0.0197).

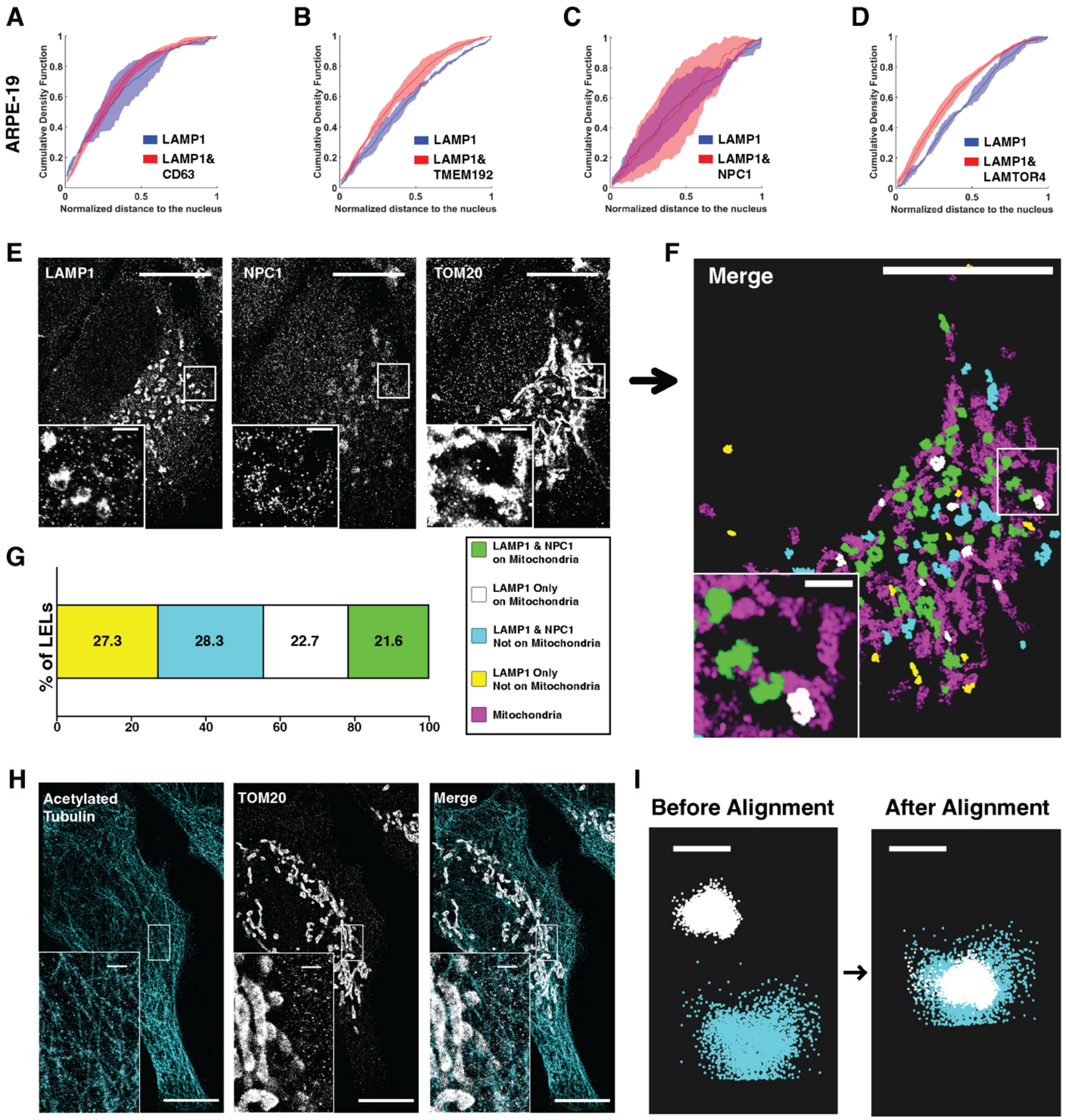


**Supplementary Figure 5: Examining LEL subpopulations with additional cellular context in ARPE-19 cells and LAMP1-GFP overexpressing HeLa cells, and multiplexed imaging controls.**

**A-D:** Cumulative density function plots of LEL distance from the nucleus from dual-color DNA-PAINT imaging experiments in ARPE-19 cells for subpopulations containing LAMP1 only or LAMP1 and target protein. Distance to the nucleus was normalized per cell to the maximum distance from the nucleus of an LEL in that cell, with values closest to zero indicating greatest proximity to the nucleus. Line indicates median with standard deviation between biological replicates. Kolmogorov-Smirnov tests were performed on the mean distributions from 3 independent biological replicates: for CD63 n = 13 cells, P = 0.193, no significance; for TMEM192 n = 14 cells, P = 0.261, no significance; for NPC1 n = 14 cells, P =0.140, no significance; for LAMTOR4 n = 14 cells, P = 0.193, no significance.

**E-G:** Representative 3-color DNA-PAINT image of LAMP1, NPC1, and mitochondria (TOM20) in ARPE-19 cells. Raw images **(G)** and post-processed image **(H)** showing a spatial map of LELs with or without NPC1 in relation to mitochondria. Quantification **(I)** shows combined subpopulations of 7 cells from 3 biological replicates. Cell scale bars, 10 µm. Inset scale bars, 1 µm.

**H:** Acetylated tubulin and mitochondria (TOM20) in HeLa cells imaged using two rabbit primary antibodies pre-labeled with secondary rabbit nanobodies, showing minimal cross-talk between the two targets. Cell scale bars, 10 µm. Inset scale bars, 1 µm.

**I:** Localizations corresponding to Tetraspeck beads imaged in a multicolor DNA-PAINT acquisition were used for image registration. Localizations are shown before and after post-processing alignment. Scale bar, 100 nm.
